# The Tandem Duplicator Phenotype is a prevalent genome-wide cancer configuration driven by distinct gene mutations

**DOI:** 10.1101/240648

**Authors:** Francesca Menghi, Floris P Barthel, Vinod Yadav, Ming Tang, Bo Ji, Zhonghui Tang, Gregory W. Carter, Yijun Ruan, Ralph Scully, Roel G. W. Verhaak, Jos Jonkers, Edison T. Liu

**Affiliations:** The Jackson Laboratory for Genomic Medicine, Farmington, CT 06030, USA; MD Anderson Cancer Center, Houston, TX 77030, USA; The Jackson Laboratory, Bar Harbor, ME 04609, USA; Division of Hematology Oncology, Department of Medicine, and Cancer Research Institute, Beth Israel Deaconess Medical Center and Harvard Medical School, Boston, MA 02215, USA; Netherlands Cancer Institute, Amsterdam, 1066 CX, Netherlands

## Abstract

The tandem duplicator phenotype (TDP) is a genome-wide instability configuration primarily observed in breast, ovarian and endometrial carcinomas. Here, we stratify TDP tumors by classifying their tandem duplications (TDs) into three span intervals, with modal values of 11 Kb, 231 Kb, and 1.7 Mb. TDPs with prominent ~11 Kb TDs feature the conjoint loss of *TP53* and *BRCA1*. TDPs with ~231 Kb and ~1.7 Mb TDs associate with CCNE1 pathway activation or *CDK12* disruptions, in conjunction with *TP53* mutations. We prove the driver role of *TP53* and *BRCA1* abrogation for TDP induction by generating short-span TDP mammary tumors in genetically modified mouse models harboring deleterious mutations in only these two genes. Lastly, heterogeneous combinations of mutations mediated by TDs are selected for and contribute to the oncogenic burden of TDP tumors.

## INTRODUCTION

Whole genome sequencing of large numbers of human cancers has revealed recurrent patterns of highly complex genomic rearrangements, such as chromothripsis, kataegis, and chromoplexy (Alexandrov et al., 2013; Baca et al., 2013; Stephens et al., 2011). The unique characteristics of these complex genomic configurations are that these structural mutations appear to have occurred simultaneously and not accreted over tumor progression. Moreover, these structural changes appear to be a strategy to equivalently alter the function of a number of oncogenic elements rather than to rely on the activation of a single driver mutation. On the other hand, as with the example of tumors characterized by high microsatellite instability, an originating mutation in a critical gene mediating genomic stability may lead to the generation of downstream genomic “scars” affecting a range of oncogenic elements that collectively drive the progression of the tumor (Kim and Park, 2014). Recently four groups have described an enrichment of head-to-tail somatic segmental tandem duplications (TDs) primarily associated with breast and ovarian cancers, which appears to follow a similar progressive mechanism and is commonly refer to as the Tandem Duplicator Phenotype (TDP) (Glodzik et al., 2017; Menghi et al., 2016; Menghi and Liu, 2016; Nik-Zainal et al., 2016; Popova et al., 2016; Watkins et al., 2016). These early reports have shown a statistical association between the TDP and loss of *BRCA1* in breast cancers (Menghi and Liu, 2016; Nik-Zainal et al., 2016), loss of *TP53* and overexpression of certain cell cycle and DNA replication genes (Menghi et al., 2016) primarily in breast and ovarian cancers, and mutations of the *CDK12* gene in a small subgroup of ovarian cancers (Popova et al., 2016). These analyses also noted that within the TDP cancer genomes, TD span sizes are clustered around specific lengths, which can be used to classify distinct genomic subtypes of TDP. In fact, we have shown that TDP tumors can be separated into at least two major subgroups: TDP group 1 tumors are BRCA1 deficient (either by epigenetic silencing or by disruptive mutations) and feature short-span TDs (~10 Kb), whereas TDP group 2 tumors are BRCA1 wild-type and feature medium-span TDs (~ 50-600 Kb) (Menghi et al., 2016; Menghi and Liu, 2016). Similarly, Nik-Zainal et al., examining over 500 breast cancer samples, described two TD-based rearrangement signatures (RS), RS1 and RS3, characterized by TDs of distinct sizes: >100 Kb (RS1) and <10 Kb (RS3) with RS3 but not RS1 strongly correlating with loss of BRCA1 (Nik-Zainal et al., 2016). Popova et al. reported the “TD plus” phenotype in some ovarian cancers featuring a large number of somatic TDs with span distribution modes at 300 Kb and 3 Mb associated with disruptive *CDK12* mutations (Popova et al., 2016). In addition, we first described the potential clinical implications of the TDP configuration with the observation that TNBC cell lines and patient derived xenografts with the TDP show significant sensitivity to cisplatin. This suggested that the underlying genomic instability for TDP, which extends beyond the “BRCAness” status, may have clinical utility.

Herein, we perform a meta-analysis of 2,717 cancer genomes representing 31 different tumor types and definitively show that the TDP is a highly recurrent chromosomal instability configuration, prevalent in triple negative breast cancer (52%), ovarian carcinoma (54%), and uterine endometrial carcinoma (48%). Only three predominant classes of TDs, characterized by discrete span size intervals centering on distribution modal values of 11 Kb, 231 Kb, and 1.7 Mb, are seen across TDP tumors, with individual genomes harboring either one single class of TDs or binary combinations of the three classes. The comprehensiveness of this study allowed us to recognize six distinct TDP subgroups classified on the basis of this robust span-size-driven TD classification. We show that the TDP subgroups occur at different frequencies across tumor types, that they are associated with specific originating genetic perturbations, and that they result in different downstream oncogenic burdens. We refine the statistical association between *TP53* and *BRCA1* mutations and short span TDP (11 Kb), and observe a strong correlation between CCNE1 pathway perturbations, *CDK12* disruptions and *TP53* mutations with the larger span TDPs (231 Kb and 1.7 Mb). Importantly, we find the identical TDP chromotype featuring predominantly short span TDs (2.5-11 Kb) in murine mammary tumors emerging only in specific genetically modified mouse models harboring the conjoint loss of the *TP53* and *BRCA1* tumor suppressor genes. The downstream gene mutations engendered by the TDs generate TDP tumors with mutational heterogeneity. Our work thus unifies a number of observations around a specific cancer genomic signature, shows how a number of genetic drivers converge on creating the TDP, and how the downstream consequences of the TDs generate oncogenic diversity that allows for further formation of genetic subgroups.

## RESULTS

### TD span distribution profiles classify TDP tumors into six distinct subgroups

Previously, using a limited number of cases, we observed that TDs found in TDP genomes fall within one of two major span size distributions with modes at ~10 Kb and ~300 Kb, respectively (Menghi et al., 2016). This is not the case for TDs found in non-TDP tumors, which typically span larger genomic regions of 1-10 Mb. This observation suggests that certain tumor types are prone to genome-wide instability manifesting in the form of DNA segmental duplications, and, given the specificity of the duplication characteristics in terms of span size, that genetic factors are driving these specific chromotypes of the TDP.

To further explore the detailed configurations of the TDP, we first analyzed the entire TCGA whole genome sequencing (WGS) dataset as the test set, comprising 25 distinct tumor types (7–59 samples per tumor type), for TD number and genomic distribution. We refer to this number and distribution as the TDP score (Menghi et al., 2016). Of the 992 TCGA cancer genomes analyzed, 118 (11.9%) were classified as TDP (**Table S1**). As predicted, we found that the TDP was not broadly distributed across cancer types, but rather was highly enriched in a subset of cancers, namely in triple negative breast cancer (TNBC), ovarian carcinoma (OV) and uterine endometrial carcinoma (UCEC), in agreement with our previous findings (**Table S1**). We then examined the TD span size distributions in individual TDP tumors and observed only a few recurrent patterns, each one characterized by either a modal or a bimodal profile (**Figure 1A**). We systematically classified these recurrent profiles by binning all of the modal peaks relative to the TD span size distributions observed across 118 identified TDP tumors in this dataset into five non-overlapping intervals, based on the best fit of a Gaussian finite mixture model (see **STAR Methods**). We then labeled the TDs corresponding to the five span size intervals as class 0: <1.6 Kb in span size; class 1: between 1.64 and 51 Kb (median value of 11 Kb); class 2: between 51 and 622 Kb (median value of 231 Kb); class 3: between 622 Kb and 6.2 Mb (median value of 1.7 Mb); and class 4: >6.2 Mb (**Figure S1**). Noticeably, classes 1-3 made up almost 95% (146/154) of all the identified modal peaks, being found in all but one TDP tumor (**Table S2**).

**Figure 1.**
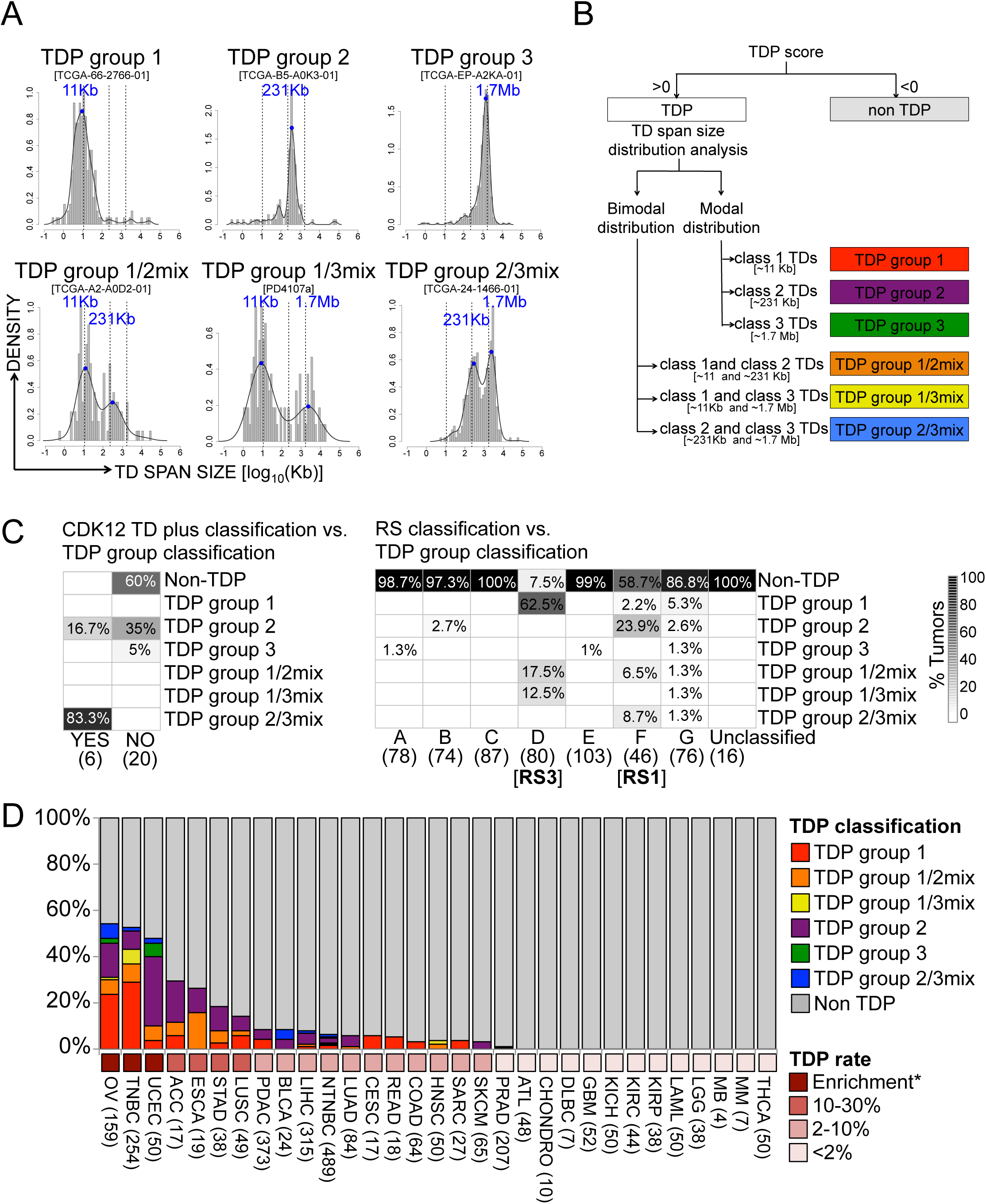
Classification of TDP genomes into six distinct subgroups. (A) Representative TD span size distribution profiles for each one of the six identified TDP subgroups. Individual distribution peaks are highlighted in blue and labeled according to the peak classification described in **Figure S1**. Also, see **Table S2**. (B) Schematic overview of the TDP subgroup classification strategy. (C) Left, The TDP group 2/3mix profile captures the large majority of tumors classified as CDK12 TD plus by Popova et al. (Popova et al., 2016). Right, altogether, the TDP groups 1, 1/2mix and1/3 mix capture over 90% of all tumors featuring RS3, according to Nik-Zainal et al. (Nik-Zainal et al., 2012). Conversely, the combination of TDP groups 2, 1/2mix and 2/3mix represent ~40% of all tumors with RS1. Numbers in parenthesis indicate the number of samples analyzed for each tumor subclass. (D) Bar chart of the relative proportion of each TDP subgroup across each one of the 31 tumor types examined. *Binomial test statistics was applied to identify tumor types that are overall enriched or depleted for the TDP. For the full list of tumor type descriptions and the exact numbers of tumor samples depicted in the bar chart refer to **Table S1**.

Using this classification, we were able to stratify TDP tumors into one of six distinct subgroups. Tumors with a modal TD span size distribution were designated as TDP group 1, group 2 or group 3, based on the presence of a single class 1 (11 Kb), class 2 (231 Kb) or class 3 (1.7 Mb) TD span size distribution peak, respectively. Tumors that showed a bimodal TD span size profile were designated as TDP group 1/2mix (featuring both a class 1 and a class 2 TD span size distribution peak), group 1/3mix (class 1 and class 3 peaks), or group 2/3mix (class 2 and class 3 peaks; **Figures 1A and B**). Only 1/118 tumors (0.8%) could not be classified into any of the six identified TDP subgroups, since it featured only very small or very large TDs (<1.6 Kb, i.e. class 0; and >6.2 Mb, i.e. class 4). This tumor was designated as ‘unclassified’ and excluded from further analysis. Thus, virtually all of the TDP tumors analyzed exhibit clearly distinct TD span size distributions converging on one of only three highly recurrent and narrowly-ranged span size intervals. These data strongly suggest that specific, distinct mechanisms of DNA instability are at play in the identified TDP subgroups.

Recently, three TD-based genomic signatures were described specifically in breast and ovarian cancers (Nik-Zainal et al., 2016; Popova et al., 2016). We asked how our classification compared with these published series. Our TDP classification algorithm classified 83% (5/6) of the reported CDK12 TD-plus phenotype-positive tumors as TDP group 2/3mix (**Figure 1C**). It also classified 93% (74/80) of RS3-positive tumors in the Sanger dataset as TDP groups 1, 1/2mix, or 1/3mix; but only 39% (18/46) of RS1-positive tumors as TDP group 2, 1/2mix or 2/3mix, with most of the remaining 61% (27/46) classifying as non-TDP (**Figure 1C**). On closer inspection, most of the tumor classified as RS1 that were not designed as TDP tumors featured only a small number of TDs (<15), which did not pass our TDP score threshold. Since our threshold was defined by a statistical segregation of a distinctive cancer genomic configuration, these subthreshold RS1-positive tumors are likely not to represent a specific mechanistic origin but a general characteristic of cancer. Thus, collectively, there is a consensus that a specific form of genomic instability characterized by accumulation of TDs exists in cancer, which we call the TDP. Our classification approach however, simplifies and unifies the identification of the TDP by generating a single score and provides refined sub-classifications based on TD span size.

### TDP subgroups occur at different frequencies across different tumor types

To confirm our classification scheme, we applied our TDP scoring algorithm to a separate dataset of whole genome sequences relative to 1725 tumor samples from individual patient donors, assembled from 30 independent studies and representing 14 different tumor types (see **STAR Methods** and **Table S3**). A total of 258 (15%) tumors from this dataset passed the threshold to be called TDP, and over 99% of these (257/258) matched one of the six TDP subgroup profiles (**Table S3**). This indicates that our classification scheme performs consistently and robustly across different tumor types and datasets.

When combined with the TCGA training set, we were able to study a total of 2717 independent tumor genomes, of which 375 (13.8%) classify as TDP (**Table S1**). Using this large dataset, we confirmed that the TDP is not a ubiquitous characteristic of cancer. In fact, whereas the TDP occurs in ~50% of TNBC, OV and UCEC tumor types, it is found in 10-30% of esophageal, gastric, squamous carcinoma of the lung, and adenoid cystic carcinomas, and in only 2-10% of a variety of other cancer types including pancreatic, liver, non-TNBC breast, and colorectal carcinomas. Finally, the TDP is highly depleted or entirely absent in leukemia, lymphoma, glioblastoma, prostate and thyroid carcinomas, and all forms of kidney cancer (**Figure 1D and Table S1**). Of note, the six TDP subgroups recurred among a few highly TDP-enriched tumor types, but at significantly different relative frequencies (**Figure 1D**). Whereas the TDP is found in almost half of all TNBC, ovarian, and UCEC tumors (52.8%, 54.1% and 48%, respectively), TDP group 1 accounts for 29% (74/254) of all TNBC tumors, and 24% (38/159) of ovarian cancers, but only for 4% (2/50) of UCEC tumors. Conversely, 30% of UCEC but only 7% of TNBC tumors, and 15% of ovarian cancers classify as the TDP group 2 (**Figure 1D and Table S1**). Taken together, these observations suggest that there must exist certain defined molecular differences that guide the formation of the distinct TDP subtypes.

### Joint abrogation of both BRCA1 and TP53 specifically drives the emergence of the TDP group 1 configuration

When we looked for specific gene mutations that may distinguish the different TDP profiles, we found several configurations that were statistically correlated with the different TDP subgroups. The most prevalent was the observation that TDP subgroups characterized by prominent short span TDs (class 1, ~11 Kb), either alone (i.e. TDP group 1) or in combination with larger TDs (i.e. TDP groups 1/2mix and 1/3mix), were tightly associated with *BRCA1* gene deficiencies and the mechanisms of disruption were either by somatic (8.4%) or germline mutation (48.7%), promoter hyper-methylation (42%), or structural rearrangement (0.9%) (**Figure 2A**). Indeed, when considering our pan-cancer dataset, <2% of non-TDP tumors showed BRCA1 deficiencies, compared with 80.9% of TDP group 1 (odds ratio [OR] = 341.4, p<2.2E-16; Fisher’s test), 60% of TDP group 1/2mix (OR = 121.2, p<2.2E-16) and TDP group 1/3mix tumors (OR = 779.5, p<2.2E-16). Importantly, this association was even stronger when analyzing the TNBC and OV datasets individually, where BRCA1 abrogation was present in at least 75% and up to 100% of TDP-group 1, TDP-group 1/2mix and TDP-group1/3mix tumors (**Figure 2A** and **Table S4**). By contrast, less than 10.5% of non-TDP and TDP-group 2 or TDP-group 3 TNBC and OV cancers share any of the mutational events affecting the *BRCA1* gene.

**Figure 2.**
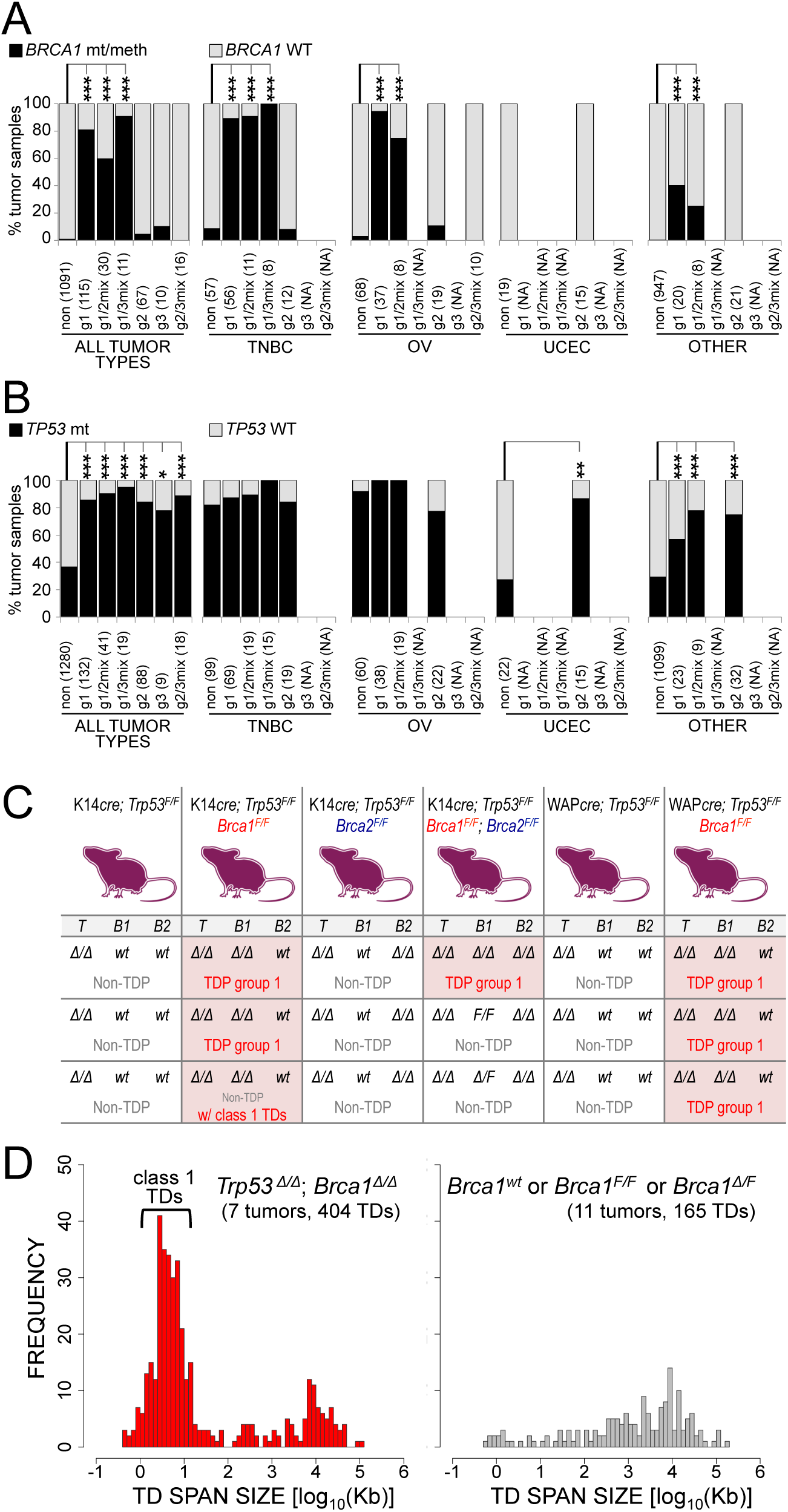
Conjoint abrogation of *BRCA1* and *TP53* results in TDP with class 1 TDs. (A) Percentage of tumor samples featuring abrogation of the *BRCA1* gene via germline or somatic mutation, promoter methylation or gene rearrangement. Only tumor type/TDP group combinations comprising at least eight tumor samples are analyzed. P-values according to Fisher’s exact test (***: p<0.001; **: p<0.01; *: p<0.05). NA: data not available; non: non-TDP tumors; g1, g1/2mix, g1/3mix, g2, g3, g2/3mix: TDP groups 1, 1/2mix, 1/3mix, 2, 3 and 2/3mix. (B) Percentage of tumor samples featuring abrogation of the TP53 gene by somatic mutation. Only tumor type/TDP group combinations comprising at least eight tumor samples are analyzed. Annotations as in (A). Number of samples for each tumor type/TDP group combination do not necessarily match those reported in (A) because of missing values. See **Table S4** for the raw data used to generate the barplots. (C) Genomic analysis of *Trp53^Δ/Δ^; Brca1^Δ/Δ^* mouse tumors: mouse models of breast cancer with somatic loss of both *Trp53* and *Brca1*, but not *Brca2* feature genome-wide scattered somatic tandem duplications whose number, distribution and span size phenocopy the TDP group 1 configuration observed in BRCA1-deficient human cancers. *T, Trp53; B1, Brca1; B2, Brca2*. Also see **Table S5**. (D) Span sizes of TDs found in *Trp53/Brca1* null tumors (left) and in Brca1-proficient tumors (right).

This indicates that BRCA1 deficiency is highly enriched in the TDP profile comprising predominantly short-span TDs (~11 Kb), either alone or in combination with larger TDs. By contrast, *BRCA2* gene disruptions are not statistically linked to any TDP configurations, either in breast and ovarian cancers or in the pan-cancer datasets (**Figure S2A**). Moreover, we found *BRCA2* mutations to be significantly depleted from the major TDP subgroups in the ovarian cancer dataset (TDP group 1 vs. non-TDP: OR = 0, p = 2.4E-03 and TDP group 2 vs. non-TDP: OR = 0, p = 3.4E-02; Fisher’s test) suggesting an exclusion of BRCA2 deficiency in TDP (**Figure S2A** and **Table S4**). These observations are in agreement with our previous finding of decreased *BRCA1*, but not *BRCA2*, expression levels in TDP tumors (Menghi et al., 2016). Taken together, these data suggest that BRCA1 deficiency by any form may be involved in the genesis of TDP configuration with a significant short-span TD population (~11 Kb), and that BRCA2 deficiency has little or no role in the TDP.

When considering the entire pan-cancer dataset, we observed a second highly prevalent mutation associated with TDP: the *TP53* gene featured significantly higher rates of somatic mutations in all TDP groups vs. non-TDP tumors (86.3% mutation rate in TDP vs. 36.7% in non-TDP, OR=10.9, p<2.2E-16; **Figure S2B**) and across each distinct TDP subgroup when compared to non-TDP tumors (TDP group 1: 85.6% mutation rate, OR=4.9, p = 5.7E-11; TDP group 2: 84.1%, OR=4.6, p = 1.4E-06; TDP group 3: 77.8%, OR=4.1, p = ns; TDP group 1/2mix: 90.2%, OR=15.2, p = 1.4E-06; TDP group 1/3mix: 94.7%, OR=19.3, p = 6.1E-05; TDP group 2/3mix: 88.9%, OR=9.0, p = 7.6E-04; Fisher’s test; **Figure 2B** and **Table S4**). This statistical association could not be found when analyzing the TNBC and OV datasets separately only because the *TP53* gene is mutated in virtually 100% of TNBC (194/226; **Table S4**) and ovarian carcinoma genomes (138/140; **Table S4**). However, a strong association between functional loss of *TP53* and TDP status was observed in the UCEC dataset, where >85% of TDP group 2 tumors share a somatic mutation of the *TP53* gene compared with <28% of non-TDP tumors (OR = 15.7, p = 6.3E-04; Fisher’s test; **Figure 2B** and **Table S4**). Taken together, these data indicate that *TP53* mutations are necessary but not sufficient for the development of all forms of TDP-related genomic instabilities. Importantly, the conjoint abrogation of both TP53 and BRCA1 was found in >72% of all breast and ovarian TDP cancers with a 11 Kb TD span peak (i.e., TDP groups 1, 1/2mix, 1/3mix), but only in <10.5% of all other TDP groups (TDP groups 2, 3 and 2/3mix) and <4.7% in non-TDP tumors (p<2.2E-16; **Figure S2C** and **Table S4**), suggesting that TDPs with predominant class 1 TDs may need the two mutations for TDP formation.

Thus far, our data suggest that loss of BRCA1 but not of BRCA2 is involved in TDP formation. Using genetically modified mouse models of mammary cancer, we sought to definitely determine the roles of TP53, BRCA1, and BRCA2 in generating the genomic pattern typical of TDP group 1. We analyzed the genomes of 18 murine mammary cancers caused by the targeted tissue-specific deletion of *Trp53* alone (n=6) or in combination with *Brca1* (n=6), *Brca2* (n=3) or both *Brca1* and *Brca2* (n=3). By assessing specific TD breakpoints, we used the identical scoring algorithm for TDP as used in human tumor samples. We found the precise configuration of TDP group 1 only in tumors with homozygous deletions of both the *Trp53* and the *Brca1* genes (**Figure 2C and Table S5**). However, there was no evidence of combined modal peaks represented by the group 1/2mix and 1/3mix configurations. Of the six tumors specifically testing the combined homozygous deletion of *Trp53* and *Brca1* showing a *Trp53^Δ/Δ^; Brca1^Δ/Δ^* genotype, five were classified as TDP group 1. Similar to the human TDP group 1 tumors, the murine mammary cancers exhibited short TD spans of 2.5-11 Kb (median value = 6.3 Kb; **Figure 2D**). The remaining *Trp53 ^Δ/Δ^*; *Brca1^Δ/Δ^* tumor that was not scored as TDP had the appropriate tandem duplication short span modal peak but did not achieve the strict numerical threshold to be called a TDP tumor (TDP score = -0.23 with cut off being 0) (Figures 2C). None of the tumors arising from sole disruption of *TP53*, or of *TP53* and *BRCA2* showed any TDP characteristics (**Figure 2C and Table S5**). In tumors arising from mice with the intention of knocking out *TP53, BRCA1*, and *BRCA2* simultaneously, we observed that whereas the *TP53* and *BRCA2* genes were affected by homozygous deletions across all three tumors, the *BRCA1* gene was found to exhibit homozygous deletion in only one tumor. Importantly, this was the only tumor among the three that classified as TDP group 1. The remaining two tumors were non-TDP and maintained either one or both functional copies of the *BRCA1* gene (Figures 2C and **Table S5**). These data provide the experimental proof that the TDP group 1 configuration is a universal and specific feature of BRCA1-linked breast tumorigenesis, emerging in the context of a *TP53* null genotype. This also implies that *BRCA1* haplo-insufficiency is not sufficient to induce the TDP in the presence of *TP53* loss, despite recent evidence that it may indeed contribute to the transformation of normal mammary epithelial cells (Pathania et al., 2011). Also, not only does *BRCA2* deficiency not induce any form of TDP, but our observations suggest that abrogation of the *BRCA2* gene does not suppress TD formation in the presence of BRCA1 deficiency. Finally, the absence of any mixed modal peak configurations (i.e., TDP groups 1/2mix or 1/3mix) suggests that additional mutations may be necessary to drive the combinatorial TDP forms.

### Identification of the genetic perturbations driving non-BRCA1-linked TDP groups

In order to identify potential genetic drivers for the non-BRCA1-linked TDPs, we compared rates of gene perturbation by somatic single nucleotide variation (SNV) across different TDP subgroups. In an initial discovery phase, we aimed at increasing the statistical power of the analysis by looking at tumor samples in the breast, ovarian and endometrial cancer datasets, which comprise the highest number of TDP tumors spread across the major subgroups, and then compared individual gene mutation rates across the TDP subgroups and in non-TDP tumors. We searched for genes whose mutation rate is significantly higher in non-BRCA1-linked TDP groups compared with TDP group 1 and with non-TDP tumors (see **STAR Methods**). *CDK12*, encoding the cyclin-dependent kinase 12, emerged as the strongest candidate gene linked to the TDP group 2/3mix profile, showing disruptive mutations in 26.7% of TDP group 2/3mix tumors, compared with 0% of TDP group 1 (p = 2.3E-04, Fisher’s test) and <1% of non-TDP tumors (odds ratio [OR] = 40.6, p = 4.0E-05, Fisher’s test; **Figure S3A**). Also, as previously reported (Popova et al., 2016), when looking at *CDK12* mutation rates within individual tumor types, the highest frequency of mutation occurred in the ovarian cancer subset, where disruption of CDK12 by somatic mutation explained 60% (6/10) of all TDP group 2/3mix tumors, but was absent in either TDP group 1 (0/27) and in non-TDP (0/45) tumors (**Figure 3A and Table S4**). Taken together, these results confirm the existence of a CDK12-linked genomic instability profile characterized by TD’s of specifically large span size (>231 Kb).

**Figure 3.**
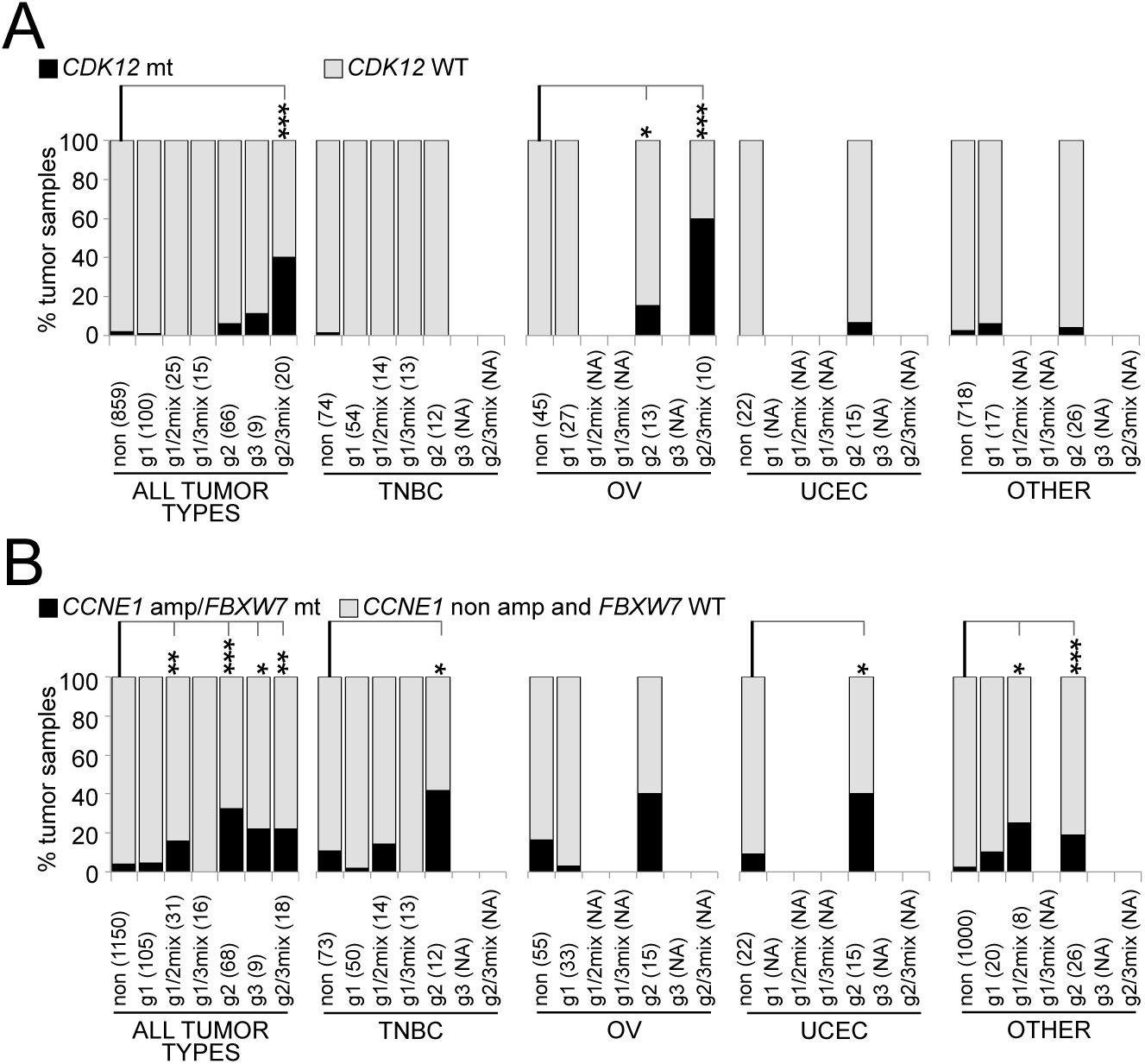
Genetic perturbations associated with BRCA1-proficient TDP groups. (A) Damaging mutations affecting the *CDK12* gene associate with TDP group 2/3mix in ovarian carcinomas (OV). Only tumor type/TDP group combinations comprising at least eight tumor samples are analyzed. P-values according to Fisher’s exact test (***: p<0.001; **: p<0.01; *: p<0.05). NA: data not available; non: non-TDP tumors; g1, g1/2mix, g1/3mix, g2, g3, g2/3mix: TDP groups 1, 1/2mix, 1/3mix, 2, 3 and 2/3mix. (B) CCNE1 pathway activation (by somatic mutation of *FBXW7* or *CCNE1* gene amplification) associates with TDP group 2 across different tumor types. Number of samples for each tumor type/TDP group combination do not necessarily match those reported in (A) because of missing values. See **Table S4** for the raw data used to generate the barplots. Also see **Figure S3**.

When focusing on TDP group 2 tumors, the strongest association involved the *FBXW7* gene, which appeared to be mutated in 11.5% of TDP group 2 tumors, compared with 2.1% of TDP group 1 (OR = 6.1, p = 2.3E-02, Fisher’s test) and 1.3% of non-TDP tumors (OR = 10, p = 4.4E-04, Fisher’s test; **Figure S3B**). Although highly significant, the disruption of *FBXW7* can only explain a modest fraction of all TDP group 2 tumors. We therefore hypothesized that other genes may be contributing to this profile by virtue of copy number variation (CNV). To explore this possibility, we focused on the TCGA dataset and examined CNV profiles that might be associated with the TDP (**Table S6**). Using a linear mixed model analysis to identify associations between TDP and gene-based CNVs while controlling for the inherent variation due to the assessment of multiple tumor types simultaneously, we ranked genes based on their significant association with TDP group 2 relative to TDP group 1 and, independently, to non-TDP tumors. The top six genes ranked in this analysis are all part of the 19q12 cancer amplicon that is frequently found in ovarian, breast and endometrial carcinomas (Etemadmoghadam et al., 2013) and that comprises the *CCNE1* and *URI1* oncogenes as well as *C19orf12, PLEKHF1, VSTM2B* and *POP4* (**Figure S3C**). The *FBXW7* gene protein product is known to act as a negative regulator of cyclin E activity by binding directly to cyclin E and targeting it for ubiquitin-mediated degradation (Klotz et al., 2009). Thus, the *FBXW7* disruptive mutations found in TDP tumors might phenocopy *CCNE1* gene amplification. The two mutations could therefore be considered as one oncogenic pathway. When assessing the frequency of CCNE1 pathway activation defined by the presence of either *FBXW7* somatic damaging mutations or *CCNE1* gene amplification, <5% of non-TDP tumors scored positively, compared with 32.4% of TDP group 2 (OR = 11.7, p = 3.5E-13), 22.2% of TDP group 3 (OR = 7, p = 4.8E-02), 16.1% of TDP group 1/2mix (OR = 4.7, p = 8.2E-03) and 22.2% of TDP group 2/3mix tumors (OR = 7, p = 5.4E-03), but only 4.8% of TDP group 1 tumors (OR = 1.2, not significant; **Figure 3B** and **Table S4**). Specifically, in each one of the individual TNBC, OV and UCEC datasets, CCNE1 pathway enhancement was found to explain at least 40% of TDP group 2 tumors (**Figure 3B**).

Taken together, though we can characterize the physical attributes of TDP into 6 structural subgroups, key genetic mutations appear to be drivers for the generation of specific TD span sizes. BRCA1 deficiency generates TDs of ~11 Kb span size, whereas CCNE1 augmentation and *CDK12* disruption are associated with TDs of specific span sizes of ~231 Kb and ~1.7 Mb. The addition of *TP53* disruption is necessary but not sufficient for both short TD and long TD configurations to emerge.

### TD breakpoint hotspots

The presence of a large number of dispersed somatic TDs affecting virtually every chromosome is the characteristic feature of TDP cancers. Nonetheless, we hypothesized that certain genomic loci may show a propensity to favor TD formation and that these loci may be different according to the class of TDs that are generated. To address this question, we identified the four major sets of TDs observed across our pan-cancer dataset (i.e. class 1 TDs (~11 Kb), class 2 TDs (~231 Kb), class 3 TDs (~1.7 Mb) and non-TDP TDs; **Figure S4A**) and for each one, we counted the number of TD breakpoints falling into consecutive 500 Kb genomic windows. We then identified genomic hotspots as 500 Kb windows with an observed number of breakpoints significantly larger than the average count value obtained from 1,000 random permutations of the TD coordinates, plus 5 standard deviations (see **STAR Methods**). A total of 245 genomic windows were identified as genomic hotspots for TD breakpoints (**Table S7**). Importantly, the overall genomic distribution of the significant hotspots appeared to be very different when comparing the four TD classes. Most of the 101 genomic hotspots relative to the non-TDP TD breakpoints tightly clustered across a small number of distinct genomic regions that have been reported to be frequently involved in oncogene amplification (i.e. *ERBB2, MYC, CCND1, CDK4* and *MDM2*, **Figures 4A**, **S4B** and **S4C**). This is in agreement with our previous report that TDs are commonly implicated in nucleating amplicon formation in regions of gene amplification in cancer (Inaki et al., 2014). Conversely, we found that the genomic hotspots corresponding to the TDP tumors were more uniformly scattered along the genome (**Figures 4B** and **S4C**) and they appeared to engage different sets of oncogenic elements, with tumor suppressor genes and oncogenes being commonly found within the genomic hotspots identified for class 1 and class 2 TDs, respectively (**Figure 4B** and see below).

**Figure 4.**
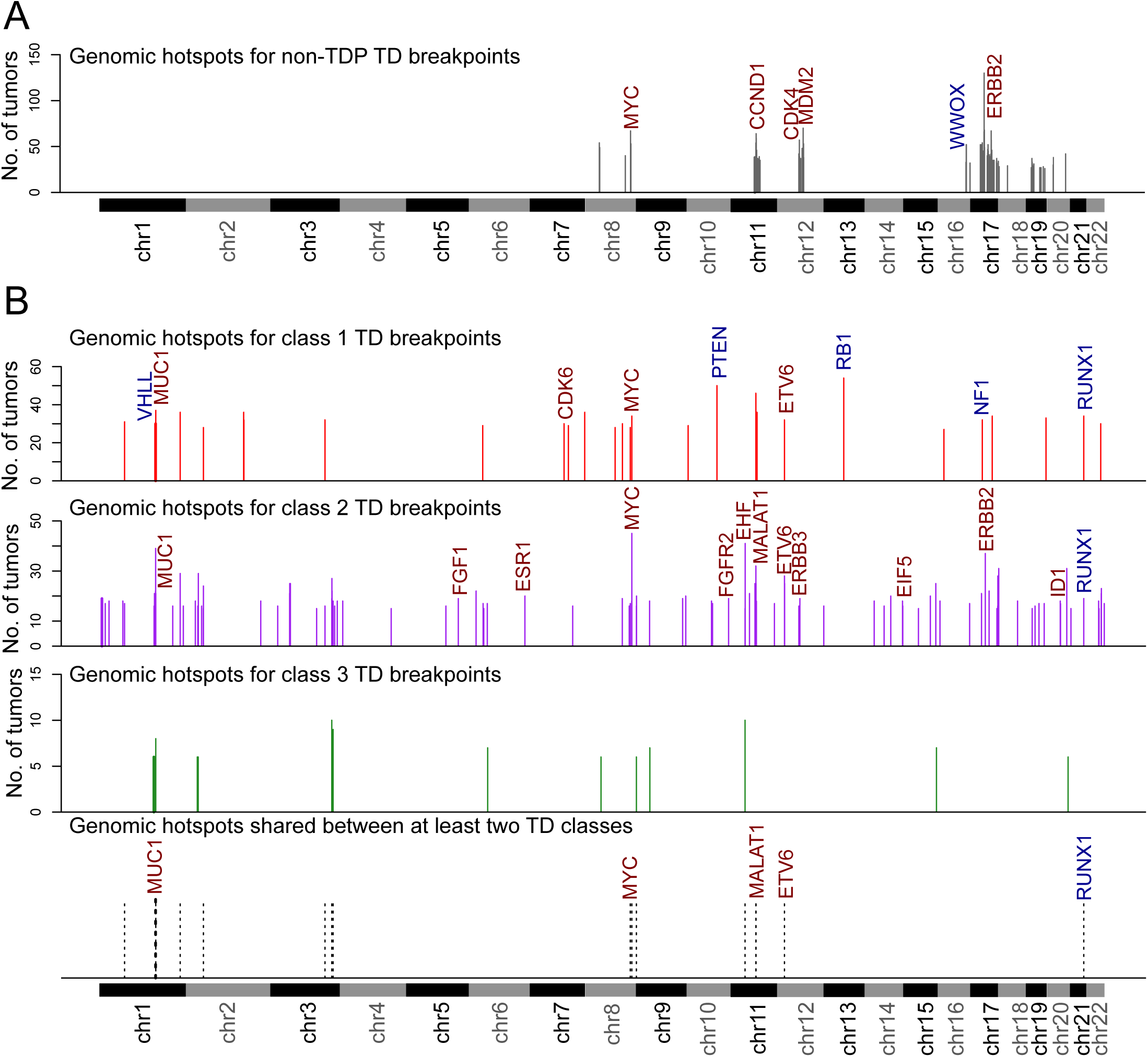
Genomic hotspots of TD breakpoints. Genomic hotspots are defined as 500 Kb windows with an observed number of TD breakpoints significantly larger than expected. (A) Genomic hotspots for TD breakpoints found in non-TDP tumors cluster at a few genomic locations, most commonly associated with oncogene amplification. (B) Genomic hotspots for TD breakpoints found in TDP tumors are more homogenously dispersed along the genome and appear to recurrently engage tumor suppressor genes (TSGs; class 1 TDs, ~11 Kb in size) or oncogenes (class 2 TDs, ~231 Kb in size). Top three panels: Genomic hotspots for class 1, class 2 and class 3 TDs. Lower panel: recurrent genomic hotspots across different TD classes. Known oncogenes and TSGs are flagged in red and blue, respectively. For the full list of the identified breakpoint hotspots refer to **Table S7**. Also see **Figure S4**.

Of note, despite the fact that the number of class 1 TDs identified in the hotspot calculation exceeded the class 2 TDs by more than two-fold (22,447 class 1 TDs vs. 9,794 class 2 TDs), there was a larger number of class 2 TD breakpoint hotspots compared with class 1 (102 vs. 30), suggesting greater selectivity for the formation of the short-span class 1 TDs (**Figure S4B** and **Table S7**).

There were several hotspots that overlapped between the TD span types accounting for 12.4% of all identified breakpoint hotspots across TDP tumors (**Figure 4B** and **Table S7**). Most of these overlaps between the class 1 and the class 2 TDs correspond to genomic regions hosting genes that may function as both oncogenes and tumor suppressor genes (TSGs) or whose activation can be accommodated by duplication or intragenic gene interruption: *ETV6* can act as both an oncogene and tumor suppressor, depending on whether it is disrupted or part of a fusion transcript; *MYC is* an oncogene that can be activated by gene duplication or by changes in the enhancer configuration; and *RUNX1* like *ETV6* acts as an oncogene with gene fusions or as a tumor suppressor. These observations suggest that the presence of primary “hotspots” for TD breakpoint formation in TDP may reflect TD proximity to cancer genes that when perturbed give a selective advantage to a tumor cell rather than any specific proclivity for instability at those sites.

### Functional consequences of TDPs: gene duplications and gene disruptions

We have previously shown that TDs occurring in the context of TDP are more likely to affect gene bodies of oncogenes and tumor suppressors than what is expected by chance alone, suggesting a strong selection for consequential genomic “scars” that favor oncogenesis (Menghi et al., 2016). Herein, we extended our analysis to account for the effect of TDs of different span sizes (class 1 vs. class 2 vs. class 3), occurring across the distinct TDP groups. A segmental duplication can affect gene body integrity in one of three ways: (1) a TD spans the entire length of a gene body resulting in gene duplication; (2) both TD breakpoints fall within the gene body resulting in a disruptive double transection; (3) only one TD breakpoint falls within a target gene body, resulting in a *de facto* gene copy number neutral rearrangement. We posited that these effects would be systematically mediated by TDs of different span sizes, with larger TDs (>231 Kb, i.e. class 2 and class 3) being mostly involved in gene duplications and shorter TDs (~11 Kb, i.e. class 1) more frequently causing gene disruptions *via* double transections. In fact, we observed that a large proportion of class 1 TDs (45%; **Figure 5A**) disrupt genes by double transection, but uncommonly result in single transections (18.2%) and even more rarely in gene duplications (5.7%), whereas the larger class 2 and class 3 TDs are more commonly implicated in single transections (66.9 and 74.7%, respectively) and in gene duplication (63.3 and 97.2%; **Figure 5A**). Importantly, these observations suggest that, by virtue of the nature of the prevalent TDs in each TDP group, distinct TDP tumor types are subjected to different forms of gene perturbation. Indeed, we found that TDP tumors featuring a prominent class 1 TD modal peak (i.e. TDP groups 1, 1/2mix, and 1/3mix) share a larger number of gene disruptions due to double transections as opposed to the other TDP tumors (p<2.2E-16; Mann–Whitney *U* test; **Figure 5B**). Conversely, TDP tumors with larger TD peaks (e.g. groups 2, 3 and 2/3mix) feature a significantly higher number of gene duplication events (p = 1.5E-10; Mann–Whitney *U* test; **Figure 5C**).

**Figure 5.**
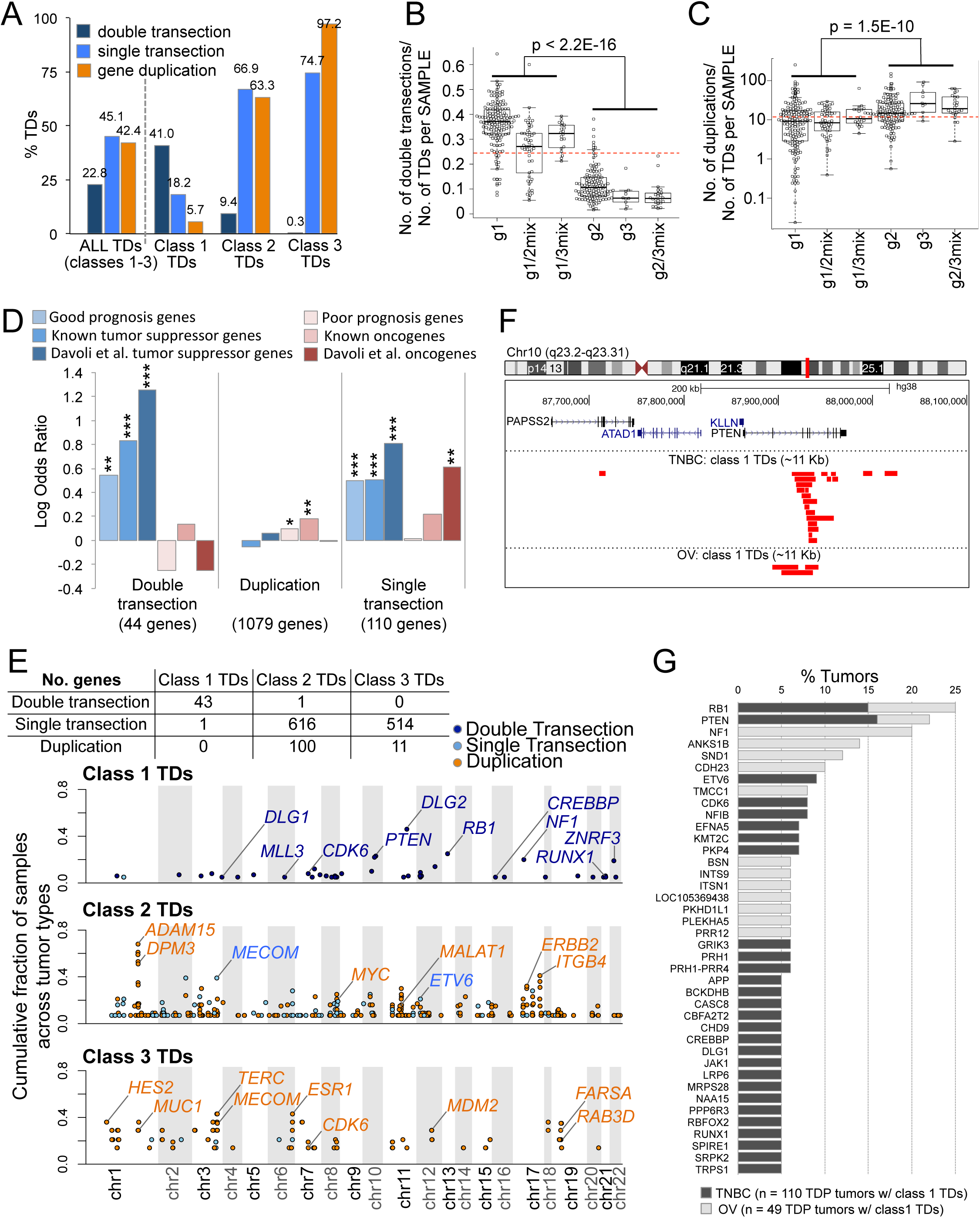
TD-mediated effects on gene bodies. (A) Different classes of TDs affect gene bodies in different ways, with shorter TDs (class 1, ~11 Kb) commonly resulting in double transection and larger TDs (class 2 and class 3, ~231 Kb and ~1.7 Mb, respectively) more frequently causing gene duplication or single transection. (B) TDP tumors featuring a prominent class 1 TD span size modal peak (TDP groups 1, 1/2mix and 1/3mix) feature a larger number of TD-mediated gene double transections compared to the other TDP tumors. P-value by Mann–Whitney *U* test. (C) TDP tumors featuring a prominent class 2 or class 3 TD span size modal peak (TDP groups 2, 3 and 2/3mix) feature a larger number of TD-mediated gene duplications compared to the other TDP tumors. P-value by Mann–Whitney *U* test. (D) Recurrently TD-impacted genes are enriched for cancer genes, with tumor suppressor genes and oncogenes being more frequently affected by double transection and duplication, respectively. TD-mediated single transections also appear to affect cancer gens more frequently than expected. P-value by Fisher’s test. (E) Recurrently TD-impacted genes by TD class and type of TD-mediated effect. The numbers of genes that are recurrently impacted by TDs found in TDP tumors, for each TD class and type of TD-mediated effect on gene body are presented in the table. The plot depicts the prevalence of these events: x_axis, genomic location; y_axis, cumulative fraction of affected TDP samples across the different tumor types examined. Selected genes are flagged for easy of reference. For tumor type specific details, see **Table S8**. (F) High density of class 1 TDs at the *PTEN* locus in both the TNBC and OV datasets. (G) Percentage of TDP tumors affected by significantly recurrent class 1 TD-mediated double transection events across the TNBC and OV datasets. Also see **Figure S5**.

Given our observation that tumor suppressor genes and oncogenes preferentially map to breakpoint hotspot regions associated with short (class 1) and larger (class 2) TDs, respectively, we predicted that these two sets of cancer genes would be directly altered by TDs in ways that augment oncogenicity. To test this hypothesis, we identified the most recurrently TD-impacted genes for each form of TD-mediated effect on gene bodies (double transection, duplication and single transection), by comparing the observed frequency and quality of TD-gene interactions with those expected based on 1,000 random permutations of the entire TD dataset (see **STAR Methods**). Using a list of tumor suppressor genes and oncogenes previously defined by three different functional criteria, we found that double transections, most commonly induced by class 1 TDs, predominantly and significantly disrupt tumors suppressor genes, whereas gene duplications, which result from class 2 and class 3 TDs, predominantly engage oncogenes but not tumor suppressors (**Figures 5D-E**). Genes undergoing single transections should theoretically result in functionally neutral copy number: one allele transected but compensated by the duplication *in situ*. However, there is primarily an enrichment of tumor suppressor genes at the sites of the single transections (**Figure 5D**). Though the precise mechanism is unclear, it is possible that the intact duplicated allele has been perturbed by either methylation, or by perturbation of the upstream regulatory mechanisms (as in topologically associating domains or TADs, see below) rendering the cell haplo-insufficient for the involved gene. Among the most commonly disrupted tumor suppressors are *PTEN* (16% and 6% of TNBC and OV TDPs with a class 1 TD span size distribution peak, i.e. TDP groups 1, 1/2mix and 1/3mix), *RB1* (15% and 10% of TNBC and OV TDPs with a class 1 TD peak), and *NF1* (affected by double transection and primarily in 20% of OV TDP tumors with a class 1 TD peak) (**Figures 5E-G** and **S5**, and **Table S8**). In the majority of the cases we examined, these highly recurrent and potentially oncogenic TD-mediated events appeared to occur independently from each other (**Tables S5A** and **S5B**). Of note, given the strong causality between loss of BRCA1 and the presence of class 1 TDs, a *BRCA1-null* status is also significantly associated with disruption of the *PTEN, RB1* and *NF1* tumor suppressor genes via TD-mediated double transection in tumor samples that are otherwise wild-type for these genes (**Figures S5A and B**). This has implications for the clinical setting since this TD-mediated suppressor gene disruption would not be detected using standard exome-sequencing protocols (discussed below).

Genes that are recurrently duplicated by TDs include: *ERBB2* (16% of UCEC, 9% of TNBC and 7% of OV TDPs with a class 2 TD span size modal peak, i.e. TDP groups 2, 1/2mix and 2/3mix); *MYC* (21% of TNBC TDP with a class 2 TD span size modal peak), and *ESR1* and *MDM2* (36% and 29%, respectively, of OV TDPs with a class 3 TD span size distribution peak, i.e. TDP groups 3, 1/3 and 2/3) (**Figures 5E** and **S5**, and **Table S8**). The oncogenic long non-coding RNA, *MALAT1*, was also often subjected to duplication in TNBC TDP tumors with class 2 modal peak (12%), suggesting its activation by gene duplication (**Figure S5A** and **Table S8**).

### Functional consequences of TDPs: duplication of regulatory elements and of chromatin structures

A recent study of breast cancer genomic rearrangements has found large span tandem duplications (>100 Kb) to frequently engage germline susceptibility loci and tissue-specific super-enhancers (Glodzik et al., 2017). Similarly, we found that cancer-associated single nucleotide polymorphisms (SNPs) identified by GWAS studies, and tissue-specific super enhancers, as a class, are indeed commonly duplicated by large span TDs, in the context of TDP tumors. In TNBCs, both class 2 and class 3 TDs are engaged in the duplication of breast-specific regulatory elements more frequently than expected, based on 1,000 permutations of TD coordinates (class 2 TDs: 17.4% observed vs. 8.7% expected, p = 3.2E-18; **Chi-squared test**; class 3 TDs: 47.6% observed vs. 35.5% expected, p =6.0E-05; **Figure 6A** and **Table S9**). Conversely, class 1 TDs are significantly less frequently involved in the duplication of these regulatory elements, even when considering their differential sequence spans (0.2% observed vs. 1.5% expected, p = 2.3E-34, **Figure 6A** and **Table S9**). Topologically associating domains (TADs) are conserved three-dimensional chromatin-folding arrangements in the genome that facilitate coordinated transcriptional regulation. Perturbations of TAD structures have been shown to be associated with transcriptional remodeling and alterations in transcriptional control (Dixon et al., 2012). This is especially true when TAD boundaries are disrupted and alternative/illigitimate enhancers are allowed to engage target gene promoters. We assessed whether TAD boundaries are disrupted by TDs in TDP tumors. Specifically, we asked whether TAD boundaries are more likely to be duplicated by a TD in breast and, independently, in ovarian cancers which are the tumor types with the largest numbers of TDP cancers. Using a 3D genome from germline cells (B lymphoblastoid cell line, GM12878) as reference (Tang et al., 2015), we mapped TD coordinates to the 3D genome. We found that TAD boundaries were statistically more frequently duplicated than expected by chance alone by class 2 TDs in both the TNBC and OV cancer datasets (TNBC: 32% observed vs. 26.3% expected, p = 2.0E-05; chi-squared test; OV: 31.6%observed vs. 26.8% expected, p = 9.36E-05; **Figure 6B** and **Table S9**). By contrast only a very modest increase in TAD boundary duplications was seen for class 3 TDs in breast cancer and no association at all was observed for class 1 TDs (**Figure 6B**). That class 2 TDs preferentially alter TAD structures can explain the observation that both tumor suppressors and oncogenes are recurrent targets of single transections mediated by these class of TDs (**Figure 5D**), since disruption of TADs via duplication of TAD boundaries may result in either a suppressive or augmenting effect on gene expression.

**Figure 6.**
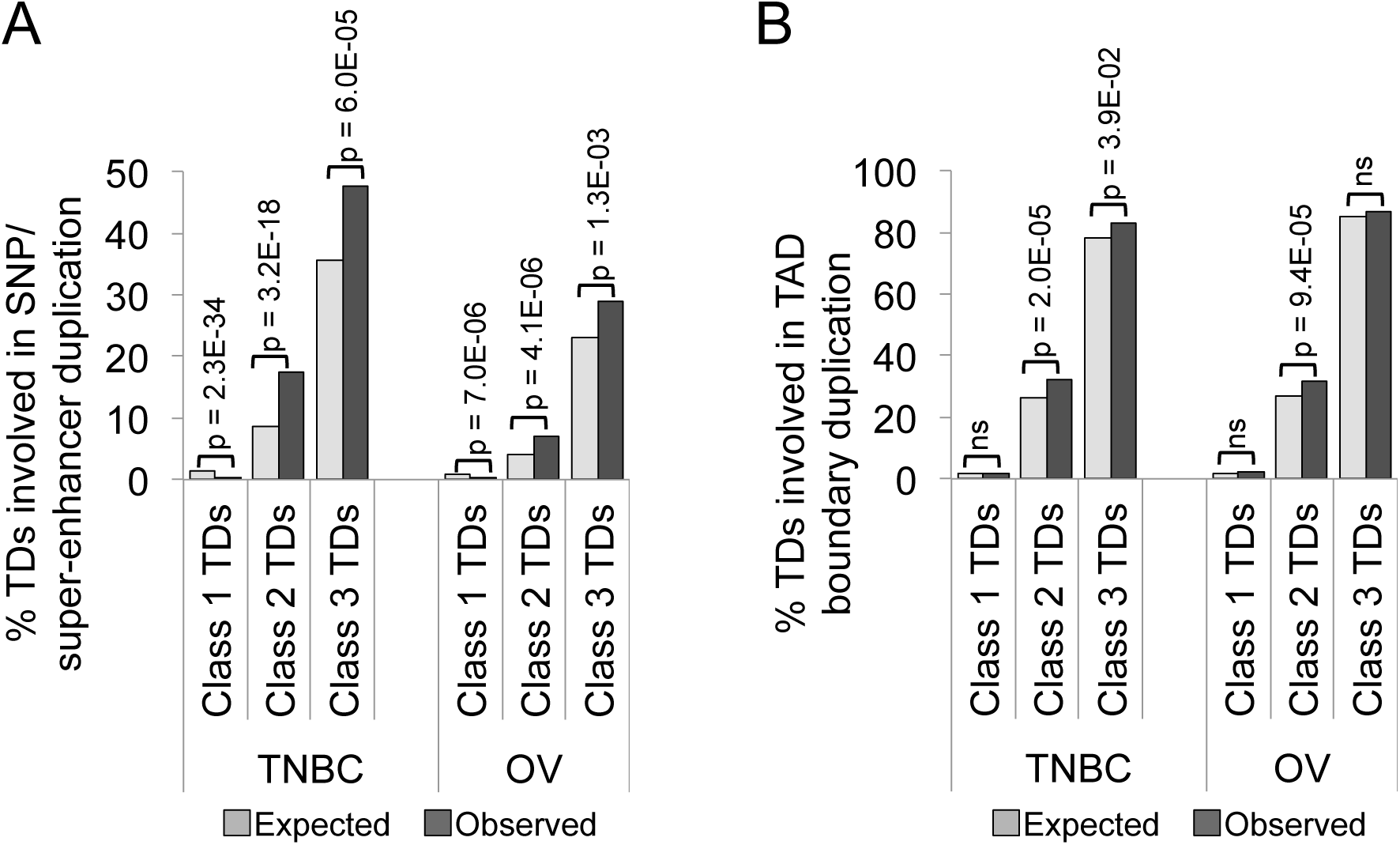
TD-mediated duplication of tissue-specific regulatory elements and TAD boundaries in TDP tumors. (A) In both the TNBC and the OV datasets, class 2 and class 3 TDs duplicate disease-associated SNPs and tissue-specific super-enhancers more frequently than what expected based on 1,000 random permutations of the TD coordinates. On the contrary, class 1 TDs less frequently engage these regulatory elements than what expected. P-values by chi-squared test. (B) In both the TNBC and the OV datasets, class 2 TDs participate in TAD boundary duplication with a significantly higher frequency than what expected based on 1,000 random permutations of the TD coordinates. P-values by chi-squared test.

Taken together, these analyses show that TDs emerging in the context of the TDP target many known oncogenic elements rather than concentrating on a few recurrent genes. We therefore asked what is the extent of the combinatorial complexity induced by TDs in the different TDP subgroups. This was measured by assessing the number of known oncogenic elements (*i.e*. oncogene, tumor suppressor gene, oncogenic lncRNA) altered by TDs for each individual tumor sample. We observed that the different TDP subgroups engage different numbers of oncogenic elements. On average, class 1 TDs found in TDP group 1 tumors result in the disruption of 3.7 known tumor suppressor genes per genome but do not engage in the duplication of other oncogenic elements (**Figures 7A** and **7B**). TDP group 1/2mix and TDP group 1/3mix have on average 2.6 disrupted TSGs, and 5.6 and 11.8 duplicated oncogenes respectively (**Figures 7A** and **7B**). By contrast, TDP groups 2, 3 and 2/3mix tumors that only feature larger span TDs rarely feature double transection of TSGs (on average 0.4, 0 and 1 TSG is affected in TDP groups 2, 3 and 2/3mix respectively), but they feature a higher number of duplications, with an average of 6.8, 37.4 and 63 duplicated oncogenes per cancer genome, respectively (**Figures 7A** and **7B**). Thus, TD formation in TDP cancer alters many oncogenic elements. The genomics “scars” in the tumors analyzed accumulate in a non-random fashion, reflecting positive selection of a range of oncogenic elements during evolution of the TDP cancer.

**Figure 7.**
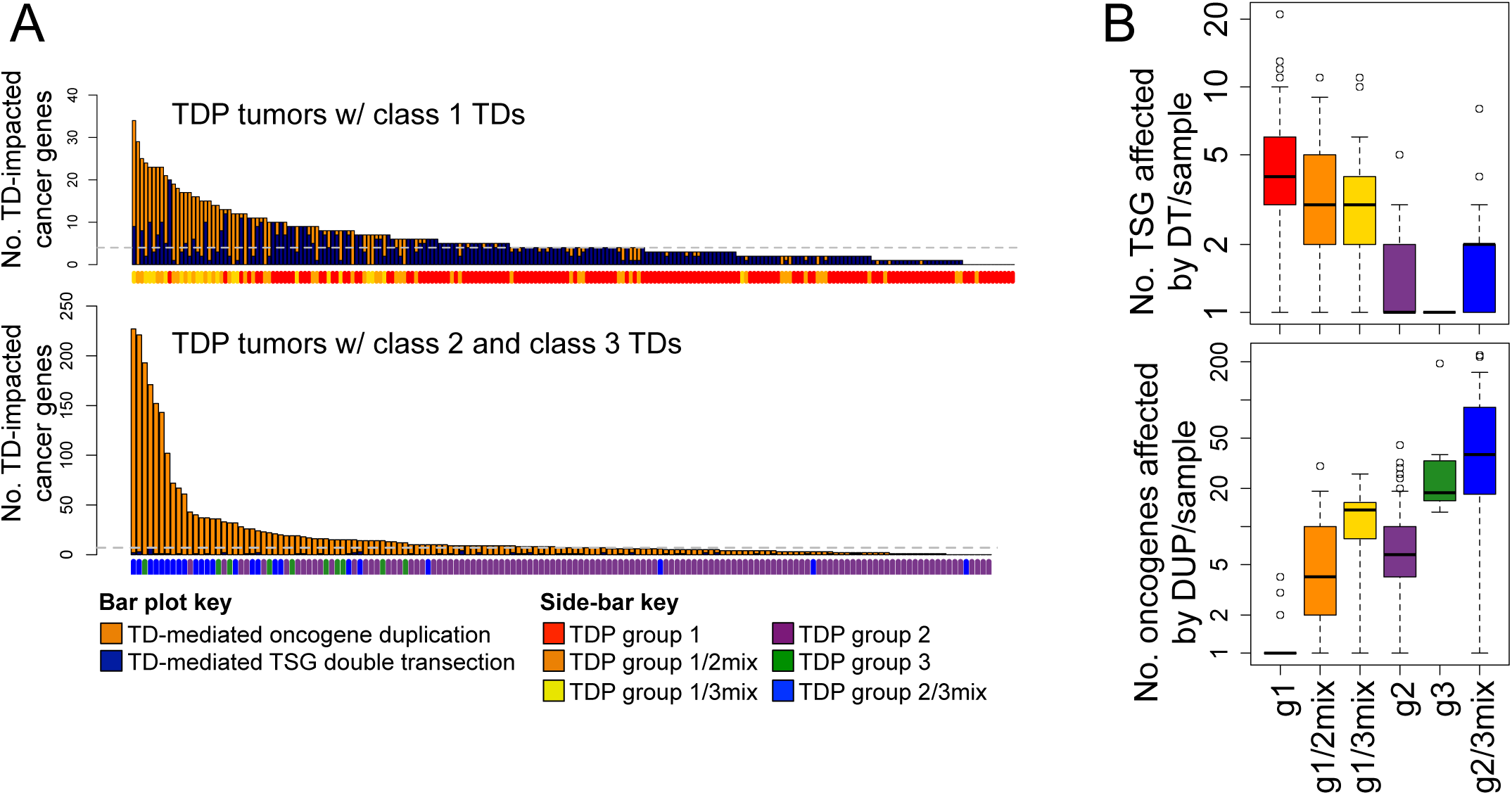
Number of TD-mediated TSG disruptions and oncogene duplications across different TDP groups. (A) Bar plots showing the number of known cancer genes that are duplicated or disrupted as a result of specific TDP configurations. (B) Box plot summary of the data presented in (A). Compared with the other tumors, TDP tumors with class 1 TDs share a larger number of TSG disruptions mediated by TD-induced double transections. Conversely, TDP tumors with class 2 and class 3 TDs have a higher incidence of oncogene duplications.

## DISCUSSION

Whole genome sequencing of tumors has revealed genome scale rearrangements generated by recurrent genetic configurations. Chromothripsis is characterized by extreme fragmentation of a chromosome site with alternating copy numbers, and chromoplexy by the presence of many translocations linked in a “daisy chain’ configuration. Chromothripsis is thought to be generated by fragmentation of lagging (and often aberrant) chromosomes in micronuclei during mitosis, and chromoplexy may be related to double strand breaks at distant DNA sites brought together by transcription factors. One of the challenges in ascertaining the frequency of these complex chromosomal configurations is that often, the classification has relied on qualitative assessment by an expert observer, limiting the reliability and reproducibility of these analyses. We resolved this problem for one chromotype, the tandem duplicator phenotype, by devising a simple quantitative scoring system that we have shown to be highly reproducible (Menghi et al., 2016). Using this algorithm, we found that the TDP is a specific genomic configuration found in ~14% of all cancer genomes across different tumor types making this a common chromotype in cancer. By better defining TDP taxonomy, we determined that TDPs can be classified by the predominant span size of their TDs: 11 Kb (i.e. class 1), 231 Kb (i.e. class 2), and 1.7 Mb (i.e. class 3). This subclassification was the key to identify the primary drivers of genome-wide TD formation. Of all TDP tumors, those characterized by the prevalence of class 1 TDs, alone (i.e. TDP group 1) or in combination with other TD span sizes (i.e. TDP groups 1/2mix, and 1/3mix) were significantly enriched for the conjoint loss of the *BRCA1* and *TP53* genes. We proved the genesis of the TDP group 1 configuration in murine models of mammary cancers that underwent conditional knock out of *TP53* and *BRCA1*. Only in the presence of BRCA1 and TP53 disruption did TDP group 1 mammary cancers emerge. By contrast, none of the *TP53* and/or *BRCA2* disrupted tumors showed any TDP configurations. Intriguingly, we did not have to modulate our TDP scoring system to accommodate the murine genome. This cross-species observation suggests that perturbation of *BRCA1* has universal genome-wide effects distinct from *BRCA2*.

Further validating this model, we have recently defined the mechanism of TD formation in *BRCA1* mutant cells in a murine cell culture model (Willis et al., 2017). We find that TDs form at sites of replication fork stalling in *BRCA1* mutant cells by a mechanism that entails re-replication of kilobases-long tracts of chromosomal DNA adjacent to the site of fork stalling. This effect is also specific to *BRCA1* loss and is not a feature of *BRCA2* loss. The striking similarities between the genetic control of TD formation in this model and the induction of group 1 TDP cancer (as defined herein) strongly suggest that class 1 TDs in cancer arise by similar aberrant re-replication at stalled forks in the presence of defective BRCA1 function and is distinct from BRCA2 action. In these experiments with murine ES cells, the *TP53* gene was not disrupted, but it is known that the p53 protein in mouse ES cells does not translocate to the nucleus in response to DNA damage and do not activate a P53 dependent response (Aladjem et al., 1998). Thus, mouse ES cells are functionally deficient in p53. Precisely how loss of *BRCA1* “licenses” class 1 TD formation and why *BRCA2* does not is currently unknown. In this regard, although BRCA1 and BRCA2 have shared roles in regulating Rad51-mediated homologous recombination and at stalled forks, BRCA1 has additional functions in double strand break (DSB) repair and in stalled fork metabolism that are not shared with BRCA2 (Aladjem et al., 1998; Bunting et al., 2010; Long et al., 2014; Pathania et al., 2011; Prakash et al., 2015; Schlacher et al., 2012; Stark et al., 2004).

For TDP subgroups without a prominent class 1 TD span size distribution peak, but characterized by the presence of larger class 2 (~ 231 Kb) and/or class 3 (~1.7 Mb) TD span size distribution peaks, the genetic origins are more heterogeneous. By association, we found that augmentation of cyclin E function either through *CCNE1* gene amplification or by *FBXW7* mutation (which augments cyclin E protein stability, (Klotz et al., 2009)) in conjunction with *TP53* disruptive mutations accounted for 40% of TDP group 2 tumors across each one of the TNBC, OV and UCEC datasets, but only manifested in 10% of non-TDP and <3% TDP group 1 of these tumor types. Intriguingly, in the larger survey of all tumor types combined, CCNE1 pathway augmentation was found to be statistically associated also with TDP groups 1/2mix, 2/3mix, and 3, albeit at lower frequencies (**Figure 3B**). This suggests that CCNE1 may play a broader role in inducing other TD span sizes. Cyclin E is known to engage cyclin dependent kinases to regulate cell cycle progression. Its dysregulation causes replicative stress by slowing replication fork progression, reducing intracellular nucleotide pools, (Bester et al., 2011), and inducing cells to enter into mitosis with short incompletely replicated genomic segments (Teixeira et al., 2015). As a model of oncogene induced replicative stress, *CCNE1* overexpression in U2OS cells induced copy number alterations which were predominantly segmental duplications (Costantino et al., 2014). It could be argued that *CCNE1* amplification is a consequence of TD formation. This is unlikely since the *CCNE1* amplifications observed in TDP group 2 cancers are detected as greater than 6 copy numbers whereas TDs constituting the TDP results in a single copy increase.

Of note, when we examined the genomic data from tumors that are not TNBC, OV, and UCEC, the associations of specific TDP subgroups with BRCA1 deficiency and CCNE1 pathway activation were significant but less in degree. Whether this is due to differences in sequencing platforms or intrinsic properties of each cancer is unclear, but it is possible that other genes may be involved in generating the TDP types in other tumor types.

Somatic mutations affecting the CDK12 gene were most prevalent in ovarian cancers where they accounted for 60% of TDP group 2/3mix tumors. Indeed, we found that the presence of a *CDK12* loss-of-function mutation in the OV dataset is invariably associated with a TDP configuration (6/6 tumors, 100%) —primarily TDP group 2/3mix. Although CDK12 disruptions can be found in TDP group 2 tumors, the predominant configuration is group 2/3mix that includes TD modal peak of 3Mb, suggesting a mechanism distinct from augmented CCNE1 function. Indeed, CDK12 is an RNA polymerase II C-terminal domain (CTD) kinase that transcriptionally regulates several homologous recombination genes. Defects in CDK12 are associated with the downregulation of critical regulators of genomic stability such as *BRCA1, ATR, FANCI*, and *FANCD2* (Blazek et al., 2011; Joshi et al., 2014). That loss of CDK12 affects *BRCA1* expression but generates a TDP profile that is clearly distinct from the BRCA1-dependent TDP group 1 configuration suggests that the primary action of CDK12 is likely to be different from its effects on the *BRCA1* gene. Although we were not able to fully explain the putative origins of a significant fraction of large-span TDP tumors, that different and distinct driver mutations ultimately generate only three fundamental TD span sizes in cancer suggests that TD formation in the context of the TDP is based on a limited number of mechanisms that tightly restrict TD formation.

The TDP is a model for combinatorial genetics in cancer. By classifying the effect of TDs on gene bodies, we showed that the TDP generates a genome-scale pro-oncogenic configuration, resulting from the modulation of tens of potential oncogenic signals through the duplication of oncogenes and regulatory elements, disruption of tumor suppressor genes, activation of oncogenic lncRNAs, and disruption of TADs. These effects are mediated systematically by TDs of different span sizes, with larger TDs (class 2 and class 3, >231 Kb) being mostly involved in gene duplications and TAD disruptions and shorter TDs (class 1, ~11 Kb) more frequently causing gene disruptions *via* double transections.

The top three genes that are disrupted by class 1 TDs are *PTEN* and *RB1* in both TNBC and OV cancer types and *NF1* in the OV dataset (**Figure 5G**). The top three genes that are disrupted by class 1 TDs are *PTEN* and *RB1* in both TNBC and OV cancer types and *NF1* in the OV dataset (**Figure 5G**). The predominant functions for these genes are in cell survival and cell cycle regulation through the PI3K, E2F and RAS pathways. However recent evidence shows a role for their products in modulating genetic instability. RB1 has been reported to be essential for DNA DSB repair by canonical non-homologous end-joining (cNHEJ), a defect invoked to explain the high incidence of genomic instability in RB1-mutant cancers (Cook et al., 2015). PTEN has been considered a major factor in genome stability through its effects on maintaining centromere stability, by controlling RAD51 expression (Shen et al., 2007), and by recruitment of RAD51 through physical association of PTEN with DNA replication forks. These studies suggest a function for PTEN with RAD51 in promoting the restart at stalled replication forks (He et al., 2015). The role of NF1 in homologous repair-deficient tumors, though statistically observed, is less established. However, the C3H-Mcm4Chaos3/Chaos3 mouse model that harbors a disruption of the minichromosome maintenance 4 (Mcm4) gene (a member of the family of MCM2-7 replicative helicases) invariably results in mammary cancers with NF1 deletions and chromosomal instability (Wallace et al., 2012). Thus, the TDP groups 1, 1/2mix, and 1/3mix tumors that originate with defects in BRCA1-mediated HR mechanisms appear to compound the defect by accumulating downstream mutations that disable genes involved in chromosomal stability and DNA repair, in addition to cellular functions such as cell cycle and cellular metabolism. By contrast, the other TDP tumors (TDP groups 2, 2/3mix, and 3) tend to activate oncogenes such as *MYC* and *ERBB2*, oncogenic lncRNAs such as *MALAT1* and to disrupt TADs. This would suggest that though the genomic characteristic is TD formation, the functional consequences of TD-induced abnormalities will vary significantly between the TDP forms.

When taken together, the totality of the current data suggests a possible mechanistic scenario for TDP induction where specific defects in homologous recombination (e.g. BRCA1 or CDK12 deficiency, but not BRCA2) and excessive replicative stress (CCNE1 pathway activation) in the presence of replication fork stalling enhance TD formation. In 91% (151/166) of TDP cancers with full genomic mutational ascertainment definitively involving one of these three driver genes, we observed concomitant mutation of *TP53*, implying that defective DNA damage checkpoint control facilitated tumorigenesis, TD formation, or both. Although disruptions of each of these genes have in the past been implicated in general genomic instability, our findings reveal that these oncogenic drivers induce a much more specific pattern of structural rearrangements (i.e., the TDP) than was previously suspected.

Our earlier study using preclinical/in *vitro* models of TNBC suggested a significant preferential sensitivity of TDP TNBC cells to cisplatin making the TDP a potential biomarker for therapeutic selection (Menghi et al., 2016). However, we had not demarcated the response based on the TDP subgroups that provide a more refined view of the tumor molecular origins. Early clinical studies suggest that platinum analogs may be more efficacious in carriers of *BRCA1* mutations, and with gross genomic evidence of homologous recombination deficiency (HRD). Our TDP analysis showed that >90% TDP TNBCs with a prominent class 1 TD span size distribution peak shared a BRCA1 deficiency. Though, the TDP appears to be an excellent genetic readout of cellular BRCA1 deficiency regardless of its mutational origin, this begs the question of whether the associated cisplatin sensitivity we observed was due to the BRCA1 status or the TDP status. In a reanalysis of the *in vitro* and *in vivo* cisplatin response data (Menghi et al., 2016), we found that sensitive cell lines included TDP group 2, BRCA1-proficent cancers. Prospective clinical studies will be needed to ascertain the detailed associations of TDP subtypes and platinum sensitivity.

The analysis of the gene disruptions as a consequence of TDP raise other therapeutic possibilities. Double transections of *PTEN* are found in 16% of TNBCs with a predominant class 1 TD span size distribution peak resulting in loss of function mutations. It has been reported in a synthetic lethal screen that PTEN knock-out cells are preferentially sensitive to PARP inhibitors (Mendes-Pereira et al., 2009). This suggests that potentially, those TDPs with PTEN disruptions may have greater deficiencies in DNA repair and may be more sensitive to a range of agents that include cisplatin and PARP inhibitors. Other frequently perturbed genes may also modulate therapeutic response: RB1, MYC, NF1. The number of known cancer genes affected by TDs range from an average of ~4 (in TDP group 1) to ~60 (in TDP group 2/3mix), with no dominant combination apparent in the datasets we have analyzed. TDP is a state where the mutational combinatorics can generate a range of potential therapeutic modifiers, some of which may be exploited to enhance treatment efficacy.

Taken together, our results provide a detailed view of a specific chromosomal configuration in cancer characterized by tandem duplications that are distributed throughout the genome that unifies a number of reports focused on individual cancers. We show, in the predominant form of TDP with class 1 short span TDs, that conjoint *BRCA1* and *TP53* mutations are essential to forming the precise TDP state. Further studies should further delineate the mechanisms of the other forms of TDP formation, and answer why these TDs are restricted to specific size ranges.

## STAR METHODS

### CONTACT FOR REAGENTAND RESOURCE SHARING

Further information and requests for resources and software should be directed to and will be fulfilled by the Lead Contact, Edison T. Liu.

## METHOD DETAILS

### Data collection for TDP classification

A catalogue of somatic tandem duplications (TDs) in human cancer was compiled from a number of published studies and a variety of sources, including The Cancer Genome Atlas (TCGA), the International Cancer Genome Consortium (ICGC) and the Catalogue Of Somatic Mutations In Cancer (COSMIC). In cases where data from two or more tumor samples from the same patient donor was available, only one sample was selected for analysis. Priority was granted to primary tumors and tumors with the highest sequence coverage. In total, 2717 tumor genomes from as many independent donors were assessed for the presence, genomic distribution and span size of somatic tandem duplications. The vast majority of the analyzed samples were primary solid tumors (n = 2,451). The dataset also included 75 metastatic solid tumors, 8 solid tumor recurrences, 18 patients derived xenografts, 55 cell lines, 98 blood tumors and 12 ascites samples (**Table S3**).

### TCGA cohort data collection and processing

Whole Genome Sequencing (WGS) data for the 992 TCGA tumors analyzed in this study has been collected from the Cancer Genomics Hub (https://cghub.ucsc.edu/). Raw reads were aligned against the reference genome *Hg19* and SpeedSeq (Chiang et al., 2015) was used to identify somatic rearrangements, as previously described (Barthel et al., 2017). Only tandem duplications with quality scores of 100 or greater and with both paired-end and split-read support were selected for TDP analysis, as these criteria have been reported to provide the highest confidence call set (Chiang et al., 2015). A list of all TCGA tumor samples analyzed with their corresponding number of somatic tandem duplications is part of **Table S3**.

### Other publicly available WGS cancer cohorts

WGS-based somatic structural variation calls from three studies (Connor et al., 2017; Ferrari et al., 2016; Fujimoto et al., 2016) were downloaded from the ICGC Data Portal (https://dcc.icgc.org/) in November 2016 (data freeze version 22). WGS-based somatic structural variation calls from 13 other studies (Bailey et al., 2016; Bass et al., 2011; Berger et al., 2011; Campbell et al., 2010; Desmedt et al., 2015; Kataoka et al., 2015; Nik-Zainal et al., 2012; Nik-Zainal et al., 2016; Northcott et al., 2012; Patch et al., 2015; Pinto et al., 2015; Stephens et al., 2009) were downloaded from the COSMIC data portal in September 2016 (data freeze version v78). Finally, WGS-based somatic structural variation calls from 13 additional independent studies were collected from the supplementary material of their corresponding publications (Baca et al., 2013; Berger et al., 2012; Grzeda et al., 2014; Hillmer et al., 2011; Imielinski et al., 2012; Inaki et al., 2014; McBride et al., 2012; Menghi et al., 2016; Natrajan et al., 2012; Ng et al., 2012; Popova et al., 2016; Totoki et al., 2014; Yang et al., 2013). A full list of all individual tumor samples collected and analyzed is reported in **Table S3**, together with annotation of their original study and WGS source.

### In-house WGS cohort and mouse tumor sequencing

Genomic libraries of 400bp size were derived from 16 PDX tumor tissues, 18 mouse tumor tissues and 2 mouse spleen tissues (normal controls) using Illumina paired end library preparation kits according to manufacturer guidelines. For the PDX samples, 150 bp paired-end sequence reads were generated using the Illumina HiSeq X Ten system and aligned to the human genome *(Hg19)*. Potential mouse contaminant reads were removed using Xenome (Conway et al., 2012). Structural variant calls were generated using four different tools (NBIC-seq (Xi et al., 2011), Crest (Wang et al., 2011), Delly (Rausch et al., 2012), and BreakDancer (Chen et al., 2009)), and high confidence events were selected when called by all four tools. In the absence of matched normal DNA samples to be used as controls, germline variants were identified as those which appear in the Database of Genomic Variants (DGV, (MacDonald et al., 2014)) and/or the 1,000 Genomes Project database (PMID:26432245). Mouse genomic libraries were sequenced using Illumina HiSeq 4000 to generate 150 bp paired-end sequence reads which were subsequently aligned to the mouse genome *(Mm10)*. Structural variants were then predicted using a custom pipeline that combines the Hydra-Multi (Lindberg et al., 2015; Malhotra et al., 2013) and SpeedSeq (Chiang et al., 2015) algorithms. Structural variation data obtained from the two spleen DNA samples were used to remove germline variants.

### The TDP classification algorithm

#### Step 1: Classification of the TCGA cohort as the test set

A TDP score was computed for each tumor sample within the TCGA cohort (n=992) based on the number and chromosomal distribution of its somatic tandem duplications (TDs), as previously described (Menghi et al., 2016). Samples with no TDs but evidence of other types of somatic rearrangements and with a minimum sequence coverage of 6X were automatically scored as non-TDP.

For each one of the 118 tumors that featured a positive TDP score, we computed the span size density distribution of all the detected TDs. Using the *turnpoints* function of the *pastecs* R package, we identified the major peak of the distribution (i.e. mode) plus any additional peaks whose density measured at least 25% of the distribution mode. A total of 154 TD span size distribution peaks were identified across the 118 TDP TCGA tumors and they appeared to cluster along recurrent and clearly distinct span-size intervals (**Figure S1**). To resolve the underlying distribution of the 154 identified TD span size distribution peaks, we used the *Mclust* function of the *mclust* R package and fit different numbers of mixture components (up to nine) to the peak distribution, using default estimates as the starting values for the iterative procedure. We compared the resulting mixture model estimates using the Bayesian information criterion and found that a mixture model comprising five Gaussian distributions with equal variance corresponded to the optimal fit. We then identified five non-overlapping span size intervals by setting thresholds corresponding to the intersections between each pair of adjacent Gaussian curves (<1.64 Kb, 1.64-51 Kb, 51-622 Kb, 622 Kb-6.2 Mb, >6.2 Mb) (**Figure S1**). Based on these thresholds, we were able to classify each TD span size distribution peak as well as each individual TD into one of 5 span size classes (classes 0-4, **Figure S1**).

Finally, we sub-grouped TDP tumors based on the presence of specific peaks/peak combinations, which appeared to be highly prevalent across the 118 TCGA TDP tumors. Tumors featuring a TD span size modal distribution were designated as TDP group 1, TDP group 2 and TDP group 3 based on the presence of a single TD span size distribution peak classified as class 1, class 2 and class 3, respectively. Similarly, tumors featuring a TD span size bimodal distribution were designated as TDP group 1/2mix (featuring class 1 and class 2 peaks), TDP group 1/3mix (featuring class 1 and class 3 peaks) and TDP group 2/3mix (featuring class 2 and class 3 peaks) (**Figure 1A** and **Table S2**). Only one out of the 118 TDP tumors did not fit any of these profiles as it featured a class 0 peak and a class 4 peak but none of the class 1, class 2 or class 3 peaks. We labeled this tumor as unclassified and did not include it in any further analysis.

#### Step 2: Validation of the TDP classification algorithm on an independent collection of sample cohorts

The TDP classification algorithm developed using the TCGA cohort as test set was applied to a completely independent dataset of 1725 tumor samples from individual patient donors, assembled from 30 different studies (referenced above) and representing 14 different tumor types. The algorithm performed consistently and robustly across the different studies of the validation cohort, by classifying 99% of the 258 TDP tumors in this cohort (257/258) into one of the six TDP subgroup profiles identified using the TCGA cohort, and by replicating similar frequencies of TDP subgroup occurrences within specific tumor types.

### SNV association analysis

Somatic single nucleotide variation (SNV) data for the tumor samples analyzed in this study was downloaded in September 2016 from the COSMIC data portal (data freeze version v78). Only tumor samples classified as breast, ovarian or endometrial carcinomas and for which whole genome or whole exome sequencing data were available were considered for the SNV-TDP group association analysis (n = 678, see **Table S3**). Only potentially damaging somatic variants were included in this analysis and comprised nonsense, frame-shift, splice site and missense mutations. Candidate genes associated with specific TDP states were considered those whose mutation rate was at least 10% and was specifically associated with only one distinct TDP profile and not any other, nor with non-TDP tumors. The significance of the associations was determined via Fisher’s exact test.

### CNV association analysis

The discovery phase of the copy number variant (CNV) association analysis was performed on the TCGA pan-cancer dataset, to allow for homogenously processed copy number information. Gene-based copy number calls relative to 977 tumor samples were obtained from the UCSC Cancer Genomic Browser (https://genome-cancer.ucsc.edu) (dataset ID: TCGA_PANCAN_gistic2, version: 2015-02-06) (**Table S3**). CNVs were classified as amplification or deletion based on GISTIC2.0 determination (Mermel et al., 2011). A liner mixed model (LMM) was used to identify the effect of TDP groups on copy number variations while controlling the variation from multiple tissues by including the tumor issue variable as random effect. Statistical analysis was performed using the package *lmerTest* (Kuznetsova, 2017) in R (version 3.3.0). P-values were adjusted for multiple testing using Benjamini-Hochberg correction. Genes were then ranked based on the p-value of their association with TDP group 2 relative to TDP group 1 and, independently, to non-TDP tumors. The top genes whose copy number change was associated with TDP group 2 tumors were identified as those with the highest cumulative rank (see also **Table S6**).

Upon identification of the 19q12 amplicon as linked to TDP group 2 status, CNV data for the *CCNE1* gene relative to the remaining tumor samples considered in this study was either retrieved from the COSMIC data portal (data freeze version v78) in the form of gene-based copy number value, or obtained from the supplementary material of the tumor samples’ original publications, when available.

### TD breakpoint analysis

Somatic TDs occurring across the entire pan-cancer dataset analyzed in this study (2717 tumor samples) were categorized into 4 classes as follows (also see **Figure S4A**):

(a) Class 1 TDs (~ 11 Kb) occurring in TDP tumors featuring a class 1 TD span size distribution peak (i.e. TDP groups 1, 1/2mix and 1/3mix; n = 22,447 TDs);
(b) Class 2 TDs (~231 Kb) from TDP tumors with a class 2 TD span size distribution peak (i.e., TDP groups 2, 1/2mix and 2/3mix; n = 9794 TDs);
(c) Class 3 TDs (~1.7 Mb) from TDP tumors with a class 3 TD span size distribution peak (i.e. TDP groups 3, 1/3mix and 2/3mix; n = 2,586 TDs) and
(d) all TDs occurring in non-TDP tumors, regardless of their individual span size (n = 25,397 TDs).

TD coordinates originally annotated using older genome assemblies were converted to the GRCh38/hg38 human genome version using the LiftOver tool of the UCSC Genome Browser (https://genome.ucsc.edu/index.html).

All of the breakpoint coordinates relative to each TD class were then binned into consecutive, non-overlapping 500 Kb genomic windows. A TD breakpoint background distribution was generated by shuffling the TD coordinates 1,000 times. At each iteration, the genomic locations of the TDs were randomly permuted across the entire genome with the exclusion of centromeric and telomeric regions, while preserving TD numbers and span sizes. Genomic hotspots for TD breakpoints were identified as 500 Kb genomic windows with an observed number of breakpoints larger than the average count value obtained from the background distribution, plus 5 standard deviations.

### Analysis of recurrently TD-impacted genes

TD-impacted genes were identified as those genes whose genomic location overlapped with that of one or more TDs. Every instance in which a gene and a TD featured some degree of genomic overlap was flagged as either (i) duplication (DUP), when the TD spanned the entire length of the gene body resulting in gene duplication; (ii) double transection (DT), when both TD breakpoints fell within the gene body resulting in the disruption of gene integrity or (iii) single transection (ST), when only one TD breakpoint fell within a target gene body, resulting in a *de facto* gene copy number neutral rearrangement. For each TD class (**Figure S4A**) and each tumor type examined, we computed the frequency with which any given gene appeared to be impacted in one of the three possible ways (i.e. DUP, DT or ST) and assigned empirical p-values to these occurrences based on the number of times, out of 1,000 iterations, that a random permutation of the TD genomic locations would result in a similar or higher frequency. Recurrently TD-impacted genes were identified as those that appeared to be affected by TDs in any one of the three possible ways in at least 5% of the tumor samples examined and in a minimum of 3 tumor samples, and with a p-value<0.05. The full list of recurrently TD-impacted genes is provided in **Table S8**.

### Cancer gene lists

***Breast cancer survival genes***. Genes associated with breast cancer patients’ prognosis data (good and poor prognosis genes) were identified as previously described (Inaki et al., 2014).

***Known cancer genes***. Lists of known tumor suppressor genes (TSGs) and oncogenes (OGs) were generated described before (Menghi et al., 2016).

***Davoli cancer genes***. Tumor suppressor genes (TSGs) and oncogenes (OGs) identified by Davoli et al. (Davoli et al., 2013).

### Analysis of disease-associated single nucleotide polymorphisms (SNPs) and tissue-specific super-enhancers

Lists of tissue-specific super-enhancers and disease-associated SNPs relative to breast and ovarian tissues were obtained from Hnisz et al. (Hnisz et al., 2013). For both tumor types examined (TNBC and OV), and for each one of the 3 major classes of TDs occurring in TDP tumors (**Figure S4A**), we computed the percentage of TDs that results in the duplication of SNPs and, separately, super-enhancers. The chi-squared test was used to compare the observed percentage to the expected one, computed as the mean value obtained from 1,000 random permutations of the TD genomic locations, as described above.

### Analysis of Topologically Associating Domains (TADs)

Genomic coordinates relative to the full catalogue of TADs for the B lymphoblastoid cell line GM12878 were published before (Tang et al., 2015). For both tumor types examined (TNBC and OV), and for each one of the 3 major classes of TDs occurring in TDP tumors (Figure S4A), we computed the percentage of TDs that overlap with TAD boundaries by at least one base pair. To compute the expected TD genomic distribution, genomic fragments were randomly sampled from non-centromere and non-telomere genomic region, with the requirement that the lengths of the sampled fragment fit the length distribution of the observed TDs. The randomly sampled fragments were then mapped to the TAD boundaries to calculate the expected percentage of TDs that overlap with TAD boundaries. The mean and standard deviation of the number of random fragments that overlap TAD boundaries were computed from 1,000 random permutations. The chi-squared test was used to compare the observed and expected values.

## QUANTIFICATION AND STATISTICAL ANALYSIS

Unless otherwise stated, statistical analysis was performed and graphics produced using the R statistical programming language version 3.3.2 (www.cran.r-project.org). All hypothesis tests were two-sided when appropriate and the precise statistical tests employed are specified in Results and figure legends.

## SUPPLEMENTAL TABLES

**Table S1, related to Figure 1D. TDP prevalence across tumor types**.

**Table S2, related to Figure 1B. TD span size distribution modal peaks across the TCGA dataset (992 tumors, 118 TDPs)**

**Table S3, related to STAR Methods. Details of the 2717 cancer genomes analyzed in this study**.

**Table S4, related to Figures 2 and 3. Statistical assessment of the putative genetic drivers for different TDP groups**.

**Table S5, related to Figure 2C. Summary Table of whole genome sequencing results for 18 mouse breast tumors**.

**Table S6, related to STAR Methods. LMM analysis of copy number data: TDP group 1 and non-TDP vs. TDP group 2**.

**Table S7, relative to Figure 4. TD breakpoint hotspots**.

**Table S8, relative to Figure 5. Recurrently TD-impacted genes**.

**Table S9, relative to Figure 6. SNP/super-enhancer and TAD boundary duplication via TDs**.

## ACKNOWLEDGMENTS

All WGS library preparation, sequencing and analysis of the mouse tumor samples were performed by JAX Cancer Center Shared Resources (Genomic Technology and the Computational Sciences) at The Jackson Laboratory for Genomic Medicine, CT 06030, USA. This work was supported by NCI grant P30CA034196 (to E.T.L.), DoD CDMRP grant BC160172 (to and E.T.L. and R.S.), the Andrea Branch and David Elliman Cancer Study Fund and the Scott R. MacKenzie Foundation (to E.T.L.), NIH grant R01CA190121 (to R.G.W.V.), CPRIT grant R140606 (to R.G.W.V.). B.J. was partially funded by the Gil Ehrich Foundation.

---

**Figure S1, related to Figure 1.**
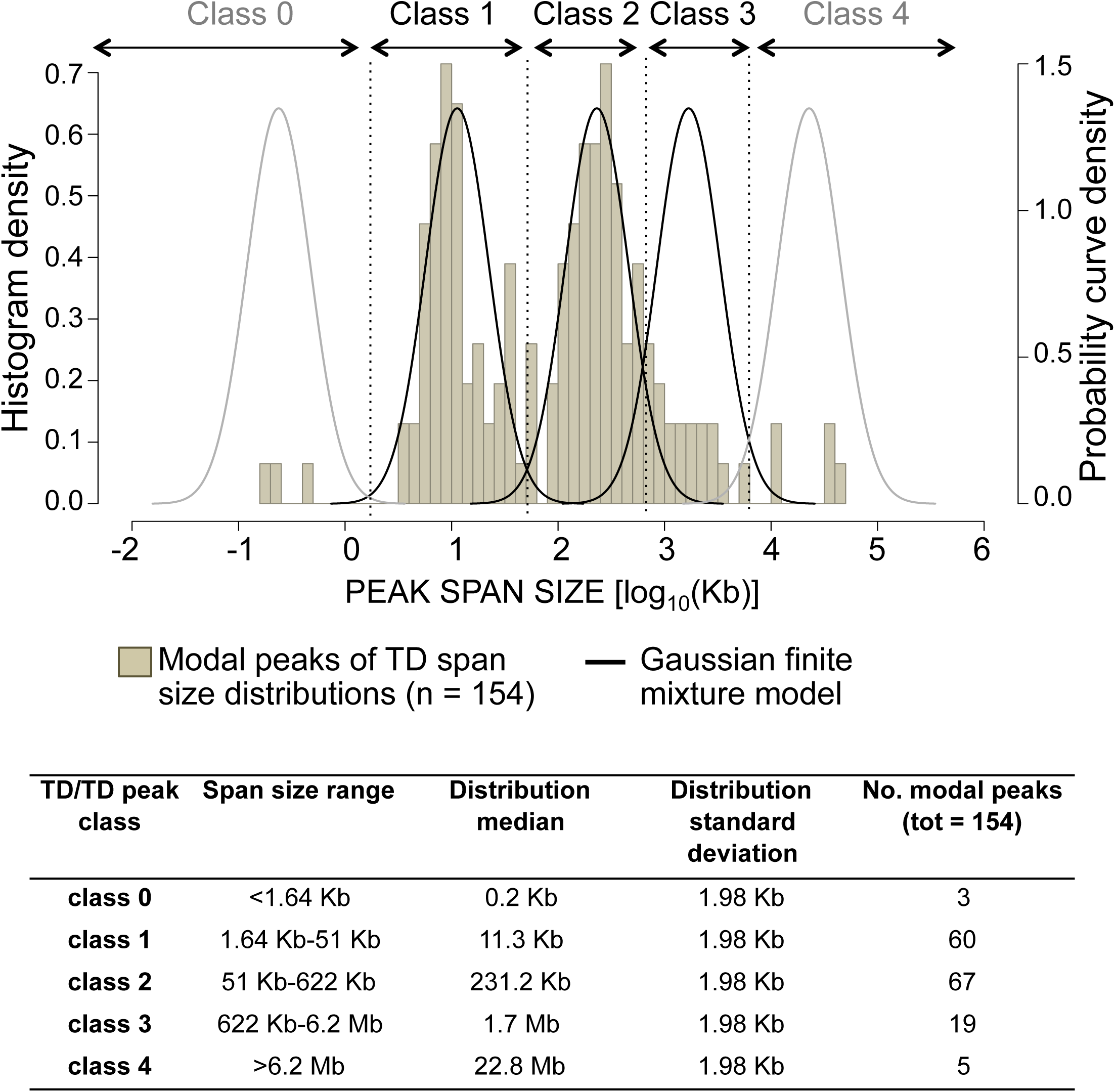
Classification of TD span size distribution peaks. Histogram: density profile of 154 TD span size distribution peaks, relative to the 118 TDP genomes included in the TCGA training pan-cancer dataset. Curves: five normal distributions sharing equal variants are identified as the best Gaussian finite mixture model to fit the TD span size peak distribution, according to the EM algorithm. Thresholds for the identification of the five TD span size intervals (i.e. classes 0–4) were set at the intersection between each pair of adjacent curves (dotted lines). Median and standard deviation values for each curve are reported in the table, together with upper and lower limits of the five identified span size intervals. Classes 1–3 comprise 146 of the 154 (94.8%) TD span size distribution peaks identified and were used to classify TDP tumors into one of six distinct TDP subgroups, as illustrated in **Figure 1B**.

**Figure S2, related to Figure 2.**
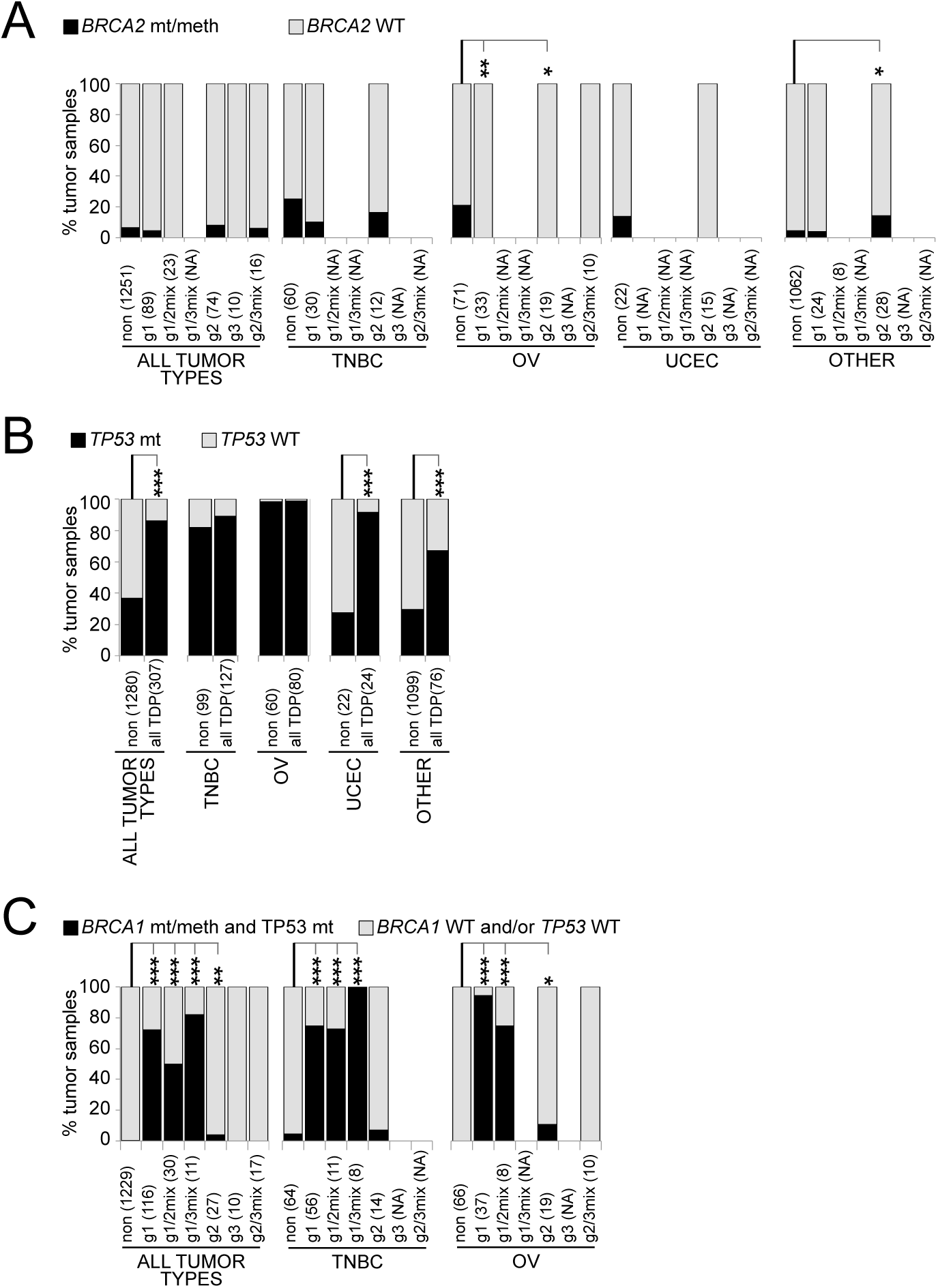
Combined abrogation of *BRCA1* and *TP53*, but not of *BRCA2* is specifically found in the large majority of TDP tumors featuring class 1 TDs. (A) Percentage of tumor samples featuring abrogation of the *BRCA2* gene via germline or somatic mutation, promoter methylation or gene rearrangement. (B) Percentage of tumor samples featuring abrogation of the TP53 gene by somatic mutation. (C) Percentage of tumor samples featuring combined abrogation of both the *BRCA1* and *TP53* genes. Only tumor type/TDP group combinations comprising at least eight tumor samples are analyzed. P-values according to Fisher’s exact test (***: p<0.001; **: p<0.01; *: p<0.05). **NA:** data not available; non: non-TDP tumors; g1, g1/2mix, g1/3mix, g2, g3, g2/3mix: TDP groups 1, 1/2mix, 1/3mix, 2, 3 and 2/3mix.

**Figure S3, related to Figure 3.**
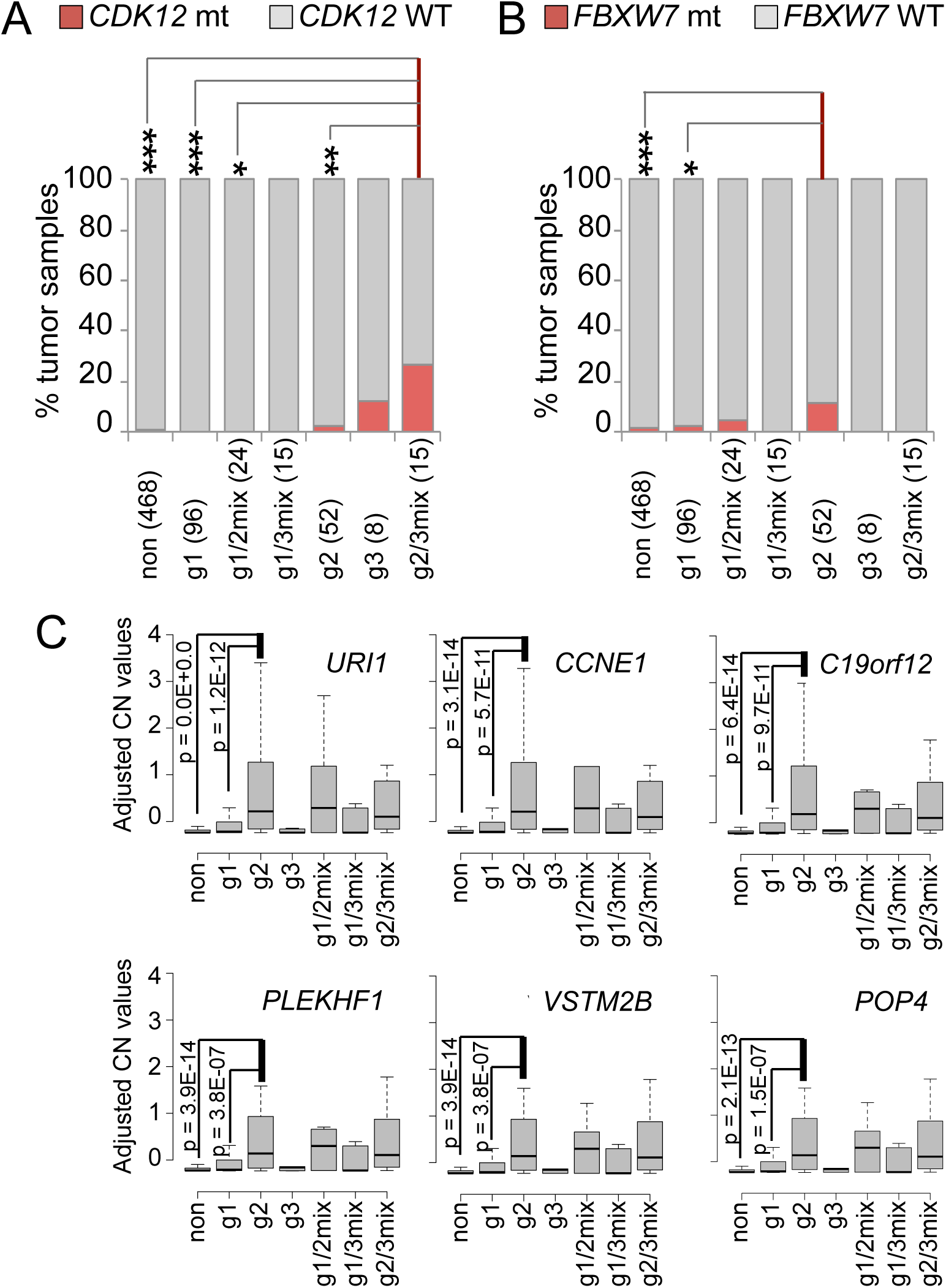
Genetic perturbations in non-BRCA1-linked TDP groups. (A, B) SNV exploratory analysis: *FBXW7* and *CDK12* mutation rates across TDP subgroups. The analysis was performed on the combined breast, ovarian and endometrial carcinoma datasets comprising a total of 678 tumors for which both somatic SNV and WGS data are available. (A) *CDK12* mutation rate. Fisher’s exact test: mix23 vs. non: C>R=40.6, P=4.0E-05; g2/3mix vs. g1: OR=inf, P=2.3E-04; g2/3mix vs. g2: OR = 17.4, p = 7.7E-03; g2/3mix vs. g1/2mix: OR = inf, p = 1.7E-02. (B) *FBXW7* gene mutation rate. Fisher’s exact test: g2 vs. non: odds ratio [OR] = 10, p = 4.4E-04; g2 vs. g1: OR = 6.1, p = 2.3E-02. (C) Chr 19q12 gene amplification in TDP group 2 tumors. Copy number distribution profile across TDP subgroups for the top six candidate genes associated with TDP group 2 in a LMM copy number analysis of the TCGA pan-cancer dataset (n = 977 tumors). All six genes map to the 19q12 amplicon and show a significant copy number increase in TDP group 2 tumors compared to non-TDP and TDP group 1 tumors. Abbreviations: non: non-TDP tumors; g1, g1/2mix, g1/3mix, g2, g3, g2/3mix: TDP groups 1, 1/2mix, 1/3mix, 2, 3 and 2/3 mix.B

**Figure S4, related to Figure 4.**
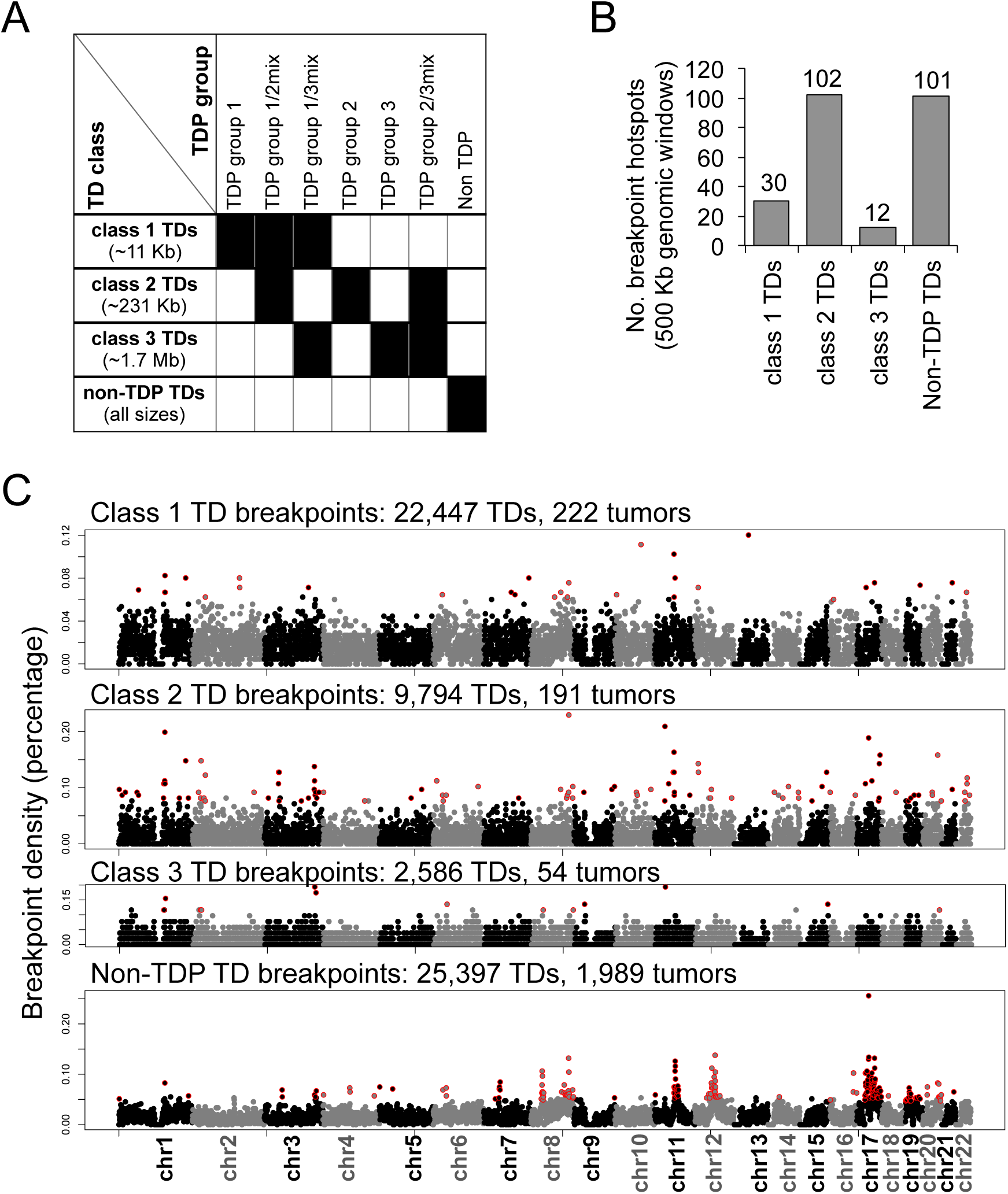
Analysis of TD breakpoint hotspots. (A) Schematic overview of the four classes of TDs considered for the breakpoint analysis. (B) Number of significant breakpoint hotspots identified across the four classes of TDs examined. Breakpoint hotpots are defined as 500 Kb genomic windows with a number of breakpoints higher than the average value obtain from 1,000 random permutation of the TD coordinates, plus 5 standard deviations. (C) TD breakpoint density per 500 Kb windows along the entire genome. Breakpoint hotspots are indicated in red and coincide with those represented in **Figure 4**. Class 2 TDs and non-TDP TDs are involved in a comparable number of breakpoint hotspots (n = 102 and 101 for class 2 and non-TDP TDs, respectively), however, their genomic distributions is remarkably different, with non-TDP TD hotspots clustering around few chromosomal locations and class 2 TD hotspots being more homogenously distributed along the entire genome.

**Figure S5, related to Figure 5.**
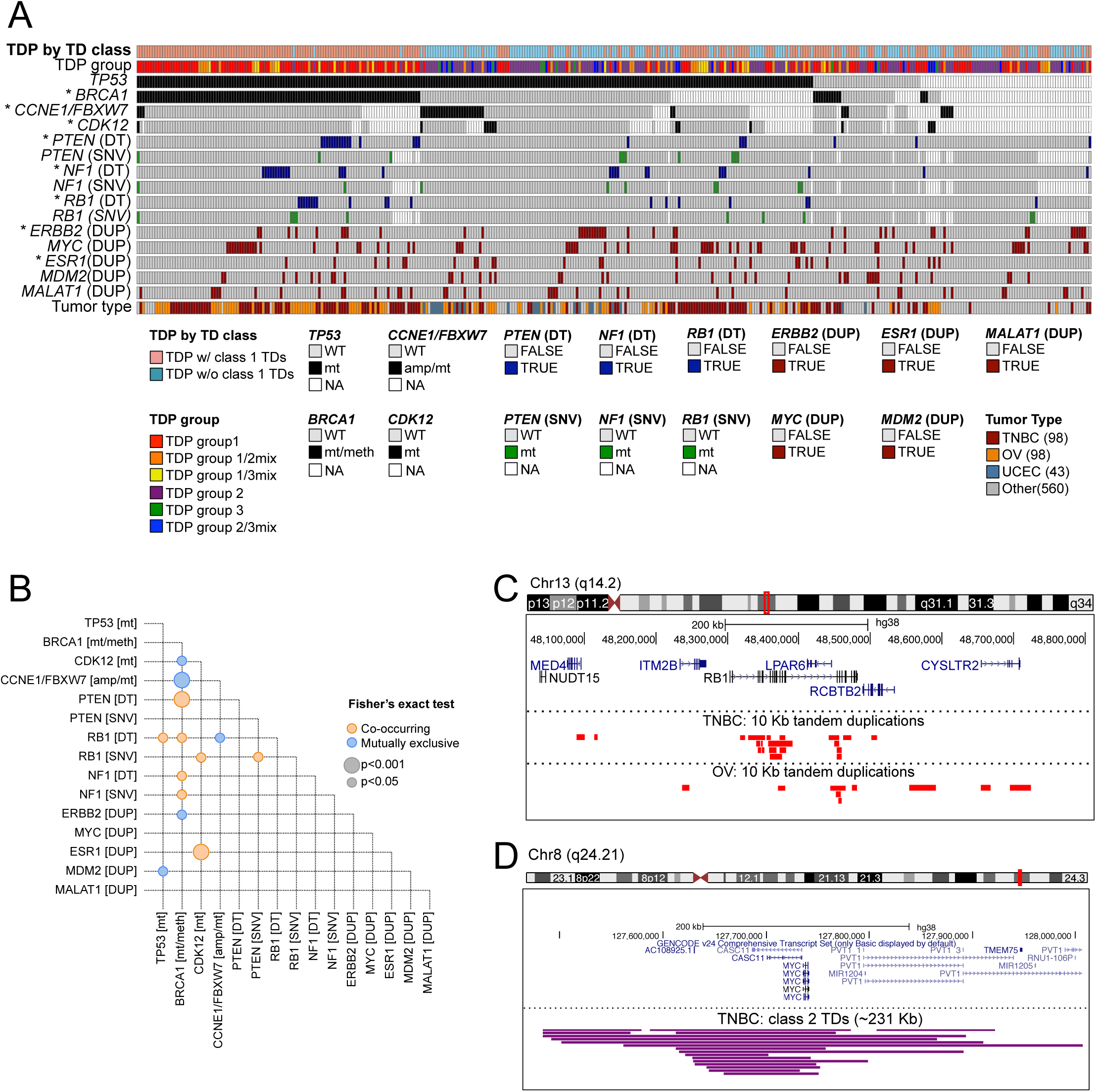
Top cancer genes most recurrently affected by TDs in TDP tumors. (A) Summary view of the major genetic perturbations and genomic consequences associated with the different TDP subgroups. Only TDP tumor are depicted in this graph. DT, double transection via TD; DUP, duplication via TD. *: variables significantly associated with TDP subgrouping (TDP by TD class) by means of Fisher's exact test. (B) Pairwise associations among TDP putative drivers and top TD-impacted genes, as depicted in (A). Only significant associations are shown (p<0.05). Orange and blue circles indicate co-occurring and mutually exclusive pairs, respectively. Among TDP putative drivers, loss of BRCA1 is mutually exclusive with activation of the CCBE1 pathway *(via CCNE1* gene amplification or *FBXW7*gene mutation) and *CDK12* gene mutation. On the other hand, loss of BRCA1 is significantly associated with disruption of the *PTEN, RB1* and *NF1* tumor suppressor genes via TD-mediated double transection in tumor samples that would otherwise be wild-type for these genes. (C) High density of class 1 TDs at the *RB1* locus in both the TNBC and OV datasets. (D) High density of class 2 TDs at the *MYC* locus in the TNBC dataset.

